# Neuronal Proteostasis is mediated by the switch-like expression of Heme-regulated Kinase 1, acting as both a sensor and effector

**DOI:** 10.1101/833178

**Authors:** Beatriz Alvarez-Castelao, Susanne tom Dieck, Claudia M. Fusco, Paul G. Donlin-Asp, Julio D. Perez, Erin M. Schuman

**Affiliations:** Synaptic Plasticity, Max Planck Institute for Brain Research, Max von Laue Strasse 4, 60438 Frankfurt am Main, Germany

## Abstract

All cells, including neurons, have regulatory feedback mechanisms that couple protein synthesis and degradation to maintain and optimize protein concentrations in the face of intra- and extracellular perturbations. We examined the feedback between the major protein degradation pathway, the ubiquitin-proteasome system (UPS), and protein synthesis in neurons. When protein degradation by the UPS was inhibited we observed a coordinate dramatic reduction in nascent protein synthesis in both neuronal cell bodies and dendrites. The mechanism for translation inhibition involved the phosphorylation of eIF2a, surprisingly mediated by eIF2a kinase 1, or heme-regulated kinase inhibitor (HRI), known for its sensitivity to heme levels in erythrocyte precursors (Han et al., 2001). Under basal conditions, neuronal expression of HRI is barely detectable. Following proteasome inhibition, HRI protein levels increase owing to stabilization of the short-lived HRI protein and enhanced translation via the increased availability of tRNAs for rare codons. Once expressed, HRI is constitutively active in neurons because endogenous heme levels are so low; HRI activity results in eIF2a phosphorylation and the resulting inhibition of translation. These data demonstrate a novel role for HRI in neurons, acting as an “immediate early protein” that senses and responds to compromised function of the proteasome to restore proteostasis.

**One sentence summary:** Proteasome inhibition leads to a compensatory reduction in neuronal protein synthesis via the stabilization and enhanced translation of short-lived HRI kinase, which is constitutively active upon expression owing to low neuronal heme levels.

## Introduction

The concept of an optimal protein concentration and its associated regulatory mechanisms is known as “proteostasis”. While many studies have focused on proteostatic regulators that are associated with the proper synthesis, folding and elimination of protein aggregation (Klaips et al., 2018), less attention has been paid to the coordination of protein synthesis and degradation within cells. The study of proteostasis is particularly important in neurons because alterations in protein synthesis and degradation, particularly via the ubiquitin proteasome system (UPS), have emerged as network hubs in many disease states (Ciechanover and Brundin, 2003; Labbadia and Morimoto, 2015; Tai and Schuman, 2008).

As an additional challenge, neurons must dynamically regulate the synaptic proteome in response to plasticity elicited by neural activity. Many studies have demonstrated that de novo protein synthesis plays an important role in synaptic transmission and plasticity, (see Sutton and Schuman, 2006; Holt et al., 2019 for review). In addition, protein degradation, mediated by the ubiquitin proteasome system (UPS, also plays an important role in regulating synaptic function (Bingol and Schuman, 2006; Tai and Schuman, 2008; Ramachandran and Margolis, 2017). In neurons, proteome regulation occurs in the cell body as well as the dendrites and axons allowing the initiation of proteome remodeling at synapses, far from the cell body (Holt et al., 2019). While it is clear that changes in synaptic transmission involve extensive regulation of the synaptic proteome via the regulated synthesis and degradation of proteins, it is not well understood how these two processes are coordinately regulated to achieve the desired level of individual proteins at synapses.

To address this question, we studied the impact of proteasome inhibition on protein synthesis in mature neurons. We found that blocking proteasome function leads to a coordinate reduction in protein synthesis, indicating the existence of a global proteostatic pathway in neurons. The entry point for down-regulation of protein synthesis is the much-studied protein synthesis initiation factor, eIF2a. The phosphorylation of eIF2a that leads to reduced neuronal translation is accomplished by one of the eIF2a kinases, heme-regulated inhibitory kinase (HRI), best known for its translational control in erythrocyte precursors. The activity and expression of neuronal HRI is regulated in a biologically clever manner, expression is kept low by a short- half-life and a translational control mechanism that is relieved by proteasome inhibition-augmenting HRI expression and leading to the coordinate regulation of protein synthesis.

## Results

### Neuronal proteasome inhibition leads to a coordinate reduction in neuronal protein synthesis, globally and locally

To determine whether the processes of neuronal protein synthesis and degradation are coupled, we examined the effect of blocking proteasomal degradation on nascent protein synthesis in cultured hippocampal neurons (Figure 1A). Blocking proteasome activity with either of two chemically distinct inhibitors (MG-132, 10 μM; Lactacystin, 10 μM; Figure 1A-E; Figure S1) for 2 hr led to a dramatic decrease in protein synthesis, measured with 2 different labeling methods: the non-canonical amino acid azidohomoalanine (AHA) and BONCAT (biorthogonal non-canonical amino acid tagging) (Dieterich et al., 2006) (Figure 1B-C) or metabolic labelling with puromycin (Figures 1D, E, and S1). The compensatory proteostatic decrease in protein synthesis did not require a complete block of proteasomal activity: dose-dependent reductions in proteasome activity led to coordinate reductions in protein synthesis (Figures 1D and S1). These data suggest that cross-talk can occur between the protein degradation and synthesis pathways in response to relatively small fluctuations in proteasome activity.

**Fig. 1.**
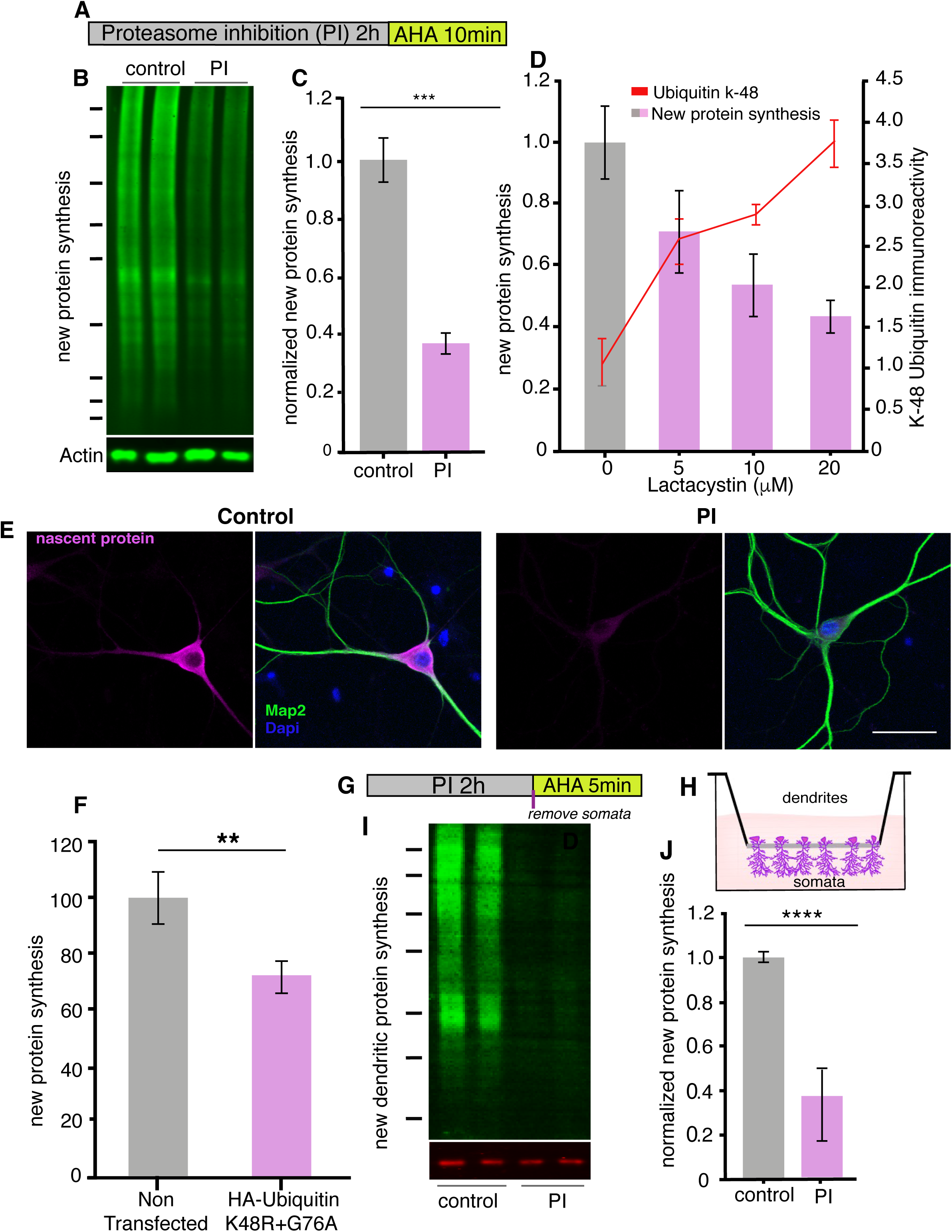
Proteasome inhibition leads to a coordinate inhibition of protein synthesis in neurons. (A) Scheme indicating experimental protocol: cultured hippocampal neurons were exposed to a proteasome inhibitor (PI; MG132 or lactacystin- see supplementary figs) for 2h and then the non-canonical amino acid AHA was added for 10 min to label newly synthesized proteins (see Methods). (B) Representative BONCAT Western blot using an anti-biotin antibody to detect newly synthesized proteins (labeled with AHA and then clicked with a biotin alkyne tag). Shown are two biological replicates each from a control sample (not treated with a PI) and a PI-treated sample, respectively. (C) Analysis of experiment shown in B. Protein synthesis in hippocampal neurons was significantly reduced following proteasome inhibition. p <0.001, unpaired t-test, 3 experiments. Error bars = SD. (D) Analysis of newly synthesized proteins (puromycylation; bar graphs) and k48 ubiquitin chains (line graph) after treatment of cultured neurons with 5, 10 or 20 μM Lactacystin. Anova (multiple comparisons) n=3 experiments for each concentration. For protein synthesis, 0 vs 5 μM p <0.001, 0 vs 10 μM p <0.01, 0 vs 20 μM p <0.001, For k48 ubiquitin, 0 vs 5 μM p <0.001, 0 vs 10 μM p <0.0001, 0 vs 20 μM p <0.001. (E) Metabolic labeling (magenta; using puromycylation, see Methods) of the global nascent protein pool in hippocampal cell bodies and dendrites following proteasome inhibition, nucleus (DAPI, blue) and dendrites (MAP2, green) can also be visualized in the images. Scale bar = 50 microns (F) Analysis of protein synthesis (using puromycylation) of the global nascent protein pool in hippocampal neurons transfected with HA-Ubiquitin K48R+G76A –see Figure S2, (unpaired t-tests p <0.001, 3 experiments, n=116 mock n=108 ubi-K48R+G76A-transfected neurons). (G) Scheme of experiment to determine whether dendritic protein synthesis is also altered, neurons were treated with PI and labeled with AHA for the last 5 minutes. (H) Scheme of the experimental set up, neurons were cultured on a special membrane that allows dendrites and axons, but not cell bodies, to grow through pores. The proteasome was inhibited, then the cell bodies were scraped away from the top of the membrane and metabolic labeling was conducted on the dendritic fraction. (I) Inhibition of the proteasome results in a coordinate inhibition of dendritic protein synthesis. Representative BONCAT Western blot showing the metabolically labeled newly synthesized dendritic protein in green. (J) Analysis of experiment shown in (I). Protein synthesis in dendrites was significantly reduced by over 60% following proteasome blockade, (p <0.05, unpaired t-test, 3 experiments) Error bars = SEM. Molecular weight markers in B and G from top-to-bottom are 250, 150, 100, 75, 50, 37, and 25 kD.

Because cultured brain preparations contain a mix of neuronal and glial cell types, we also visualized protein synthesis, *in situ*, using puromycylation to detect protein synthesis (Schmidt et al., 2009). In neurons, we observed that proteasome inhibition led to a decrease in protein synthesis apparent in both the cell bodies and dendrites of neurons (Figure 1E). The reduction in protein synthesis was not due to cell death (Figure S2) or compromised cell health as protein synthesis levels were restored following an extensive (14 hr) washout of the proteasome inhibitor (Figure S2). Furthermore, the overexpression of a mutant ubiquitin molecule, K48R+G76A, that inhibits polyubiquitination and is resistant to deubiquitinases (Hodgins et al., 1992; Ju and Xie, 2004, 2006) also decreased protein translation (Figures 1F and S2).

To determine if the proteostatic mechanism also functions in neuronal processes, we used a special membrane which separates neuronal cell bodies from dendrites and axons (“neurites”) (see Methods). We added a proteasome inhibitor for 2 hr, then physically removed the cell bodies from the region above the membrane and performed a brief (5 min) metabolic labeling (using AHA and BONCAT) on the isolated dendrites and axons inhabiting the region below the membrane (Figure 1G,H). We found that protein synthesis in neurites was also significantly inhibited by proteasome inhibition (Figure 1 I, J), indicating that the feedback mechanism coupling protein degradation to synthesis also functions locally in neuronal processes.

### Phosphorylation of the initiation factor eIF2a by HRI is responsible for reduced neuronal protein synthesis in both cell bodies and dendrites

To determine the mechanisms by which protein synthesis is reduced, we first asked whether proteasome inhibition brings about a reduction in protein synthesis by altering transcription. Blocking transcription during proteasome inhibition, however, did not affect the reduction in protein synthesis we previously observed (Figure S3). We thus focused on protein translation directly and examined the phosphorylation status, following proteasome inhibition, of several canonical translation factors known to influence protein synthesis initiation and elongation rates. The translation factors eIF4B, eIF4EBP1/2, and eIF4G showed no change in phosphorylation (Figure S3). In contrast, as has been observed in mouse embryonic fibroblasts and other cell types (Fribley et al., 2004; Jiang and Wek, 2005; Mazroui et al., 2007; Nawrocki et al., 2005; Vabulas and Hartl, 2005; Yerlikaya et al., 2008; Zhang et al., 2010), we observed a marked and rapid increase in the phosphorylation of eIF2a (a key regulator of translation initiation) (Harding et al., 2000), without a change in total eIF2a levels (Figure 2A-C). Consistent with a causal role of eIF2a phosphorylation in the inhibition of neuronal protein synthesis, we found that treatment with the small molecule inhibitor of peIF2a function ISRIB (integrated stress response inhibitor; (Sidrauski et al., 2015)) reversed the effects of proteasomal inhibition on protein synthesis (Figure 2D, E), without affecting basal levels of translation (Figure S3).

**Fig. 2.**
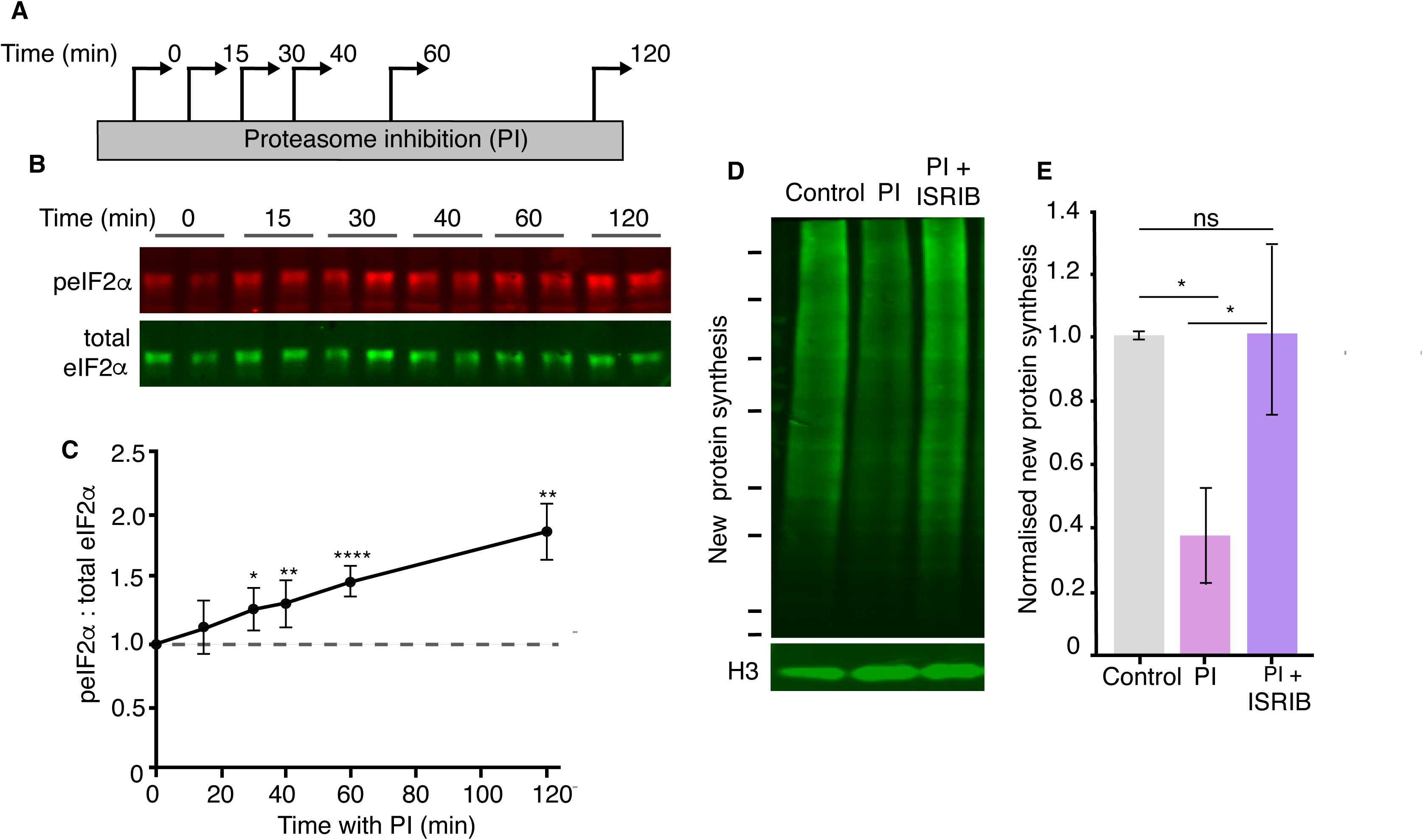
Reduced protein synthesis is mediated by eIF2a phosphorylation. (A) Scheme indicating experimental protocol: cultured hippocampal neurons were treated with a proteasome inhibitor (PI) and collected at the indicated time points. (B) Representative Western blot showing the levels of phosphorylated (detected using a phospho-specific anti eIF2a anti-body) and total eIF2a under control conditions (t = 0) or following the indicated minutes of proteasome inhibition. (C) Analysis of experiments shown in B. PI treatment led to a significant increase in peIF2a levels. (unpaired t-test control vs each time point, Control vs 15 min p<0.05, control vs 30min p>0.05, control vs 40min p≤0.01, control vs 60min p≤0.0001, control vs 120min p≤0.01. (3 experiments). Error bars = SD. (D) Representative BONCAT Western blot showing the metabolically labeled newly synthesized dendritic protein in green. Treatment with the small molecule inhibitor ISRIB rescued the PI-induced decrease in protein synthesis. Molecular weight markers from top-to-bottom are 250, 150, 100, 75, 50, 37, 25, and 20 kD. Histone H3 is shown as a loading control. (E) Analysis of experiment shown in D. The PI-induced protein synthesis inhibition (control vs. PI: p ≤0.05) was rescued by ISRIB treatment (control vs. ISRIB, p ≤0.05, unpaired t-test), 4 experiments, Error bars = SD.

There are 4 kinases that are known to phosphorylate eIF2a, eIF2a kinases 1-4, commonly known as PERK (PKR-like ER kinase), PKR (protein kinase double-stranded RNA-dependent), GCN2 (general control non-derepressible-2), and HRI (heme-regulated inhibitor) (Donnelly et al., 2013; Taniuchi et al., 2016). In order to evaluate which kinase (or kinases) phosphorylates eIF2a in response to proteasome inhibition, we first focused on GCN2 since it is well-established as a regulator of translation in neurons during synaptic plasticity (Costa-Mattioli et al., 2005; Trinh and Klann, 2013), and was previously described to be activated in response to proteasome inhibitors in *D. melanogaster* (Suraweera et al., 2012). Using cultured neurons from GCN2 knock-out mice we examined the sensitivity of protein synthesis to proteasomal inhibition. Surprisingly, in the absence of GCN2 protein synthesis was still inhibited by proteasome blockade (Figure 3A). We conducted the same experiments in cultured neurons obtained from PERK knockout mice or in PKR-inhibited neurons and again observed no effect on the proteasome-dependent inhibition of protein synthesis (Figure 3A). We thus turned our attention to the least likely candidate, HRI, a kinase that is primarily activated by reduced cellular heme levels and is known to play an important role in regulating globin translation in erythrocytes (Han et al., 2001). Using neurons from an HRI knock-out mouse (Han et al., 2001) we observed a dramatically reduced inhibition of protein synthesis induced by proteasome blockade with metabolic labeling detected by Western blot (Figure 3B,C) or *in situ* labeling of cultured hippocampal neurons (Figure 3D, E). HRI deletion had no effect on the basal levels of protein synthesis in neurons or in brain tissue (Figures 3E and S4). The absence of HRI also significantly reduced the proteasome inhibition-induced increase in eIF2a phosphorylation (Figures 3F and S4), while the absence or inhibition of the other eIF2a kinases did not (Figure S4). These data show that a kinase, known primarily for its translational regulatory role in erythrocytes, plays a critical role in neuronal proteostasis.

**Fig. 3.**
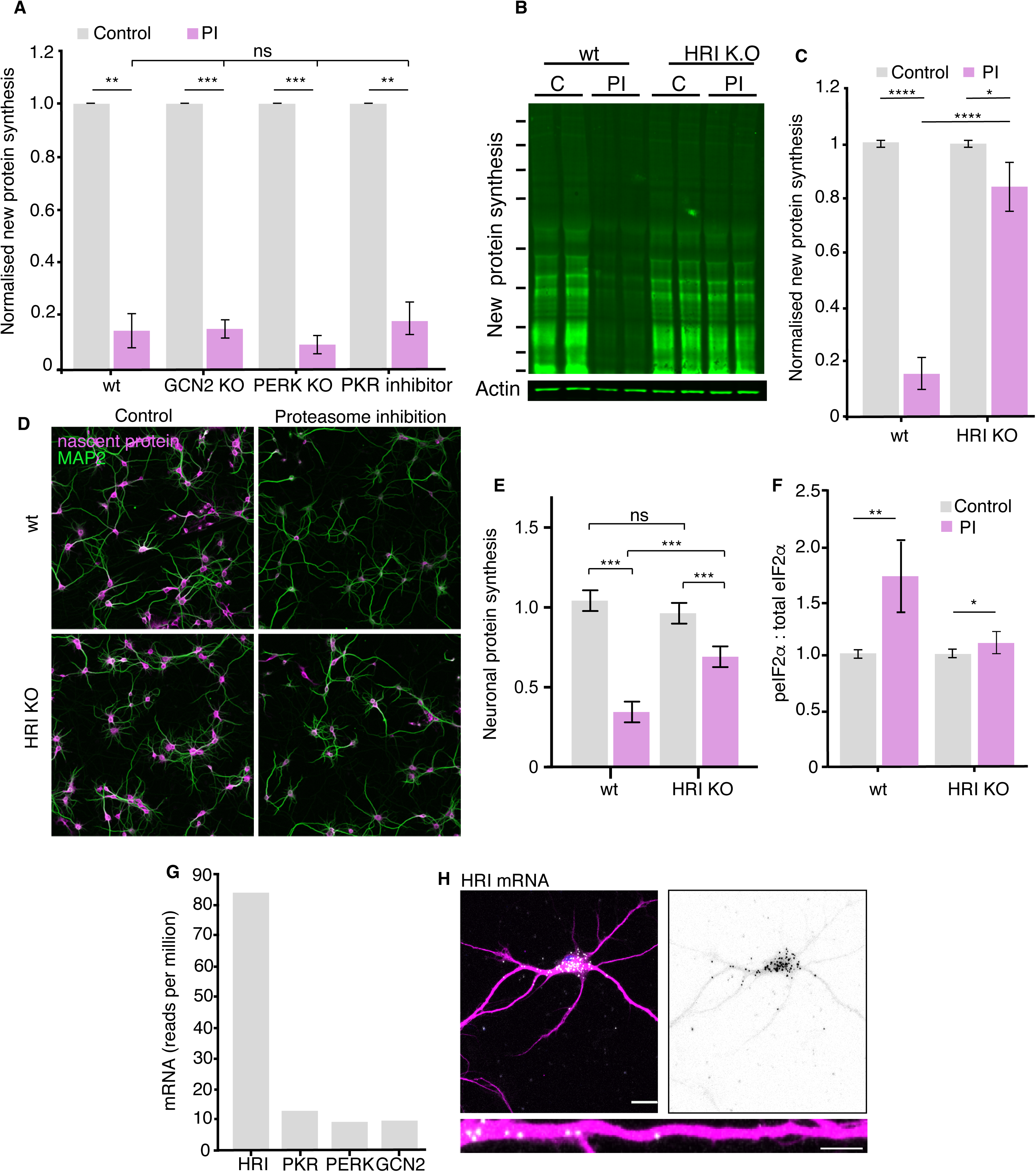
HRI kinase is responsible for the proteasome-inhibition induced increases in eIF2a phosphorylation. (A) Genetic deletion (KO) or inhibition of the eIF2a kinases GCN2, PERK or PKR did not rescue the inhibition of protein synthesis elicited by proteasome inhibition. (wt vs. kinase KO or inhibition: GCN2, PERK, unpaired t-test p ≤ 0.001 and PKR p ≤ 0.01, respectively, n = 2 experiments, 4 biological replicates per KO). Error bars = SD. (B) Representative puromycylation Western blot showing that genetic deletion of HRI kinase rescues the inhibition of protein synthesis. Molecular weight markers from top-to-bottom are 250, 150, 100, 75, 50, 37, 25, 20, and 15 kD. (C) Analysis of the experiment shown in B. The amount of PI-induced protein synthesis inhibition was significantly reduced in the HRI KO (unpaired t-test on wt PI vs KO. PI p ≤0.05, wt control vs. PI p ≤0.0001, KO. control vs. PI p ≤0.0001, n = 5 experiments, error bars SD. (D) Metabolic labeling (magenta; using puromycylation, see Methods) of the global nascent protein pool in cultured hippocampal neurons following proteasome inhibition showing that HRI KO neurons showing less PI-induced inhibition of protein synthesis. Scale bar = 100 μm. (E) Analysis of experiments like that shown in D. Protein synthesis in hippocampal neurons was significantly reduced in the HRI KO following proteasome inhibition (unpaired t-test on wt control vs. k.o. control p>0.05, wt PI vs k.o. PI p ≤0.001, wt control vs. PI p ≤ 0.001, KO control vs. PI p ≤ 0.001.n = 3 experiments, wt control n=1390, wt-PI n=955, ko control=1457, ko-PI n=992 neurons). (F) Analysis of eIF2a phosphorylation in response to proteasome inhibition. PI treatment led to a significant increase in peIF2a levels. (unpaired t-test on wt control vs. PI: p ≤ 0.01, t-test on HRI KO control vs. PI: p ≤ 0.05. 4 experiments –see Figure S4. (G) RNA-seq data from hippocampal slices indicates that HRI mRNA is the most abundant amongst the 4 eiF2a kinases (H) Representative fluorescence in situ hybridization image detecting HRI mRNA in neuronal somata and dendrites. Scale bar = 20 μm for somata and 10um for dendrites- See figure S5.

### HRI protein is expressed at vanishingly low levels in neurons under control conditions

Given the surprising involvement of HRI, and the poor characterization of its neuronal expression, we next sought to measure the expression of HRI mRNA and protein in hippocampal neurons. Using data from a recent RNA-sequencing study (Tushev et al., 2018) we found that the HRI transcript is the most abundant of the 4 eIF2A kinases (Figure 3G) in hippocampal neurons. We used fluorescence *in situ* hybridization to validate and localize HRI mRNA in hippocampal neurons and tissue, and detected it in both somata and dendrites (Figures 3H and S5). The use of HRI knock-out tissue allowed us to identify in neurons, after immunoprecipitation, an HRI-specific band with one antibody (Figure S6). We observed that the HRI expression level in brain is very low compared to the levels in iron-rich material like liver or blood (Figure S6).

We next considered whether proteasome inhibition might boost HRI expression by altering the level of HRI mRNA or protein. Quantification of HRI mRNA using droplet digital PCR (ddPCR) revealed no change in HRI mRNA levels following 2 hrs of proteasome inhibition (Figure 4A), consistent with the transcription-independence of the protein synthesis reduction elicited by proteasome inhibition (Figure S3). Following treatment with a proteasome inhibitor for 2 hrs there was, however, a significant upregulation in neuronal HRI protein levels (Figure 4B), suggesting that HRI might be a proteasome substrate. Although we were unable to detect appreciable polyubiquitylation of HRI (data not shown), using an *in vitro* assay with purified 20S proteasome and recombinant HRI we observed that HRI degraded by the proteasome (Figure S6). The observation that HRI is a proteasome substrate suggests that proteasome inhibition stabilizes HRI protein levels. To estimate the magnitude of this effect, we metabolically pulse-labeled cultured neurons with ^S35^Methionine and immunoprecipitated HRI at different chase times under control and proteasome inhibition conditions. Surprisingly, we observed that HRI protein has an extremely short half-life under control condition (∼4.2 hrs); the normal short-lived HRI protein was significantly stabilized by proteasome inhibition (Figure 4C, D). These data indicate that inhibition of the proteasome alone is sufficient to augment HRI protein levels and can account for at least part of the increase in HRI protein expression.

**Fig. 4.**
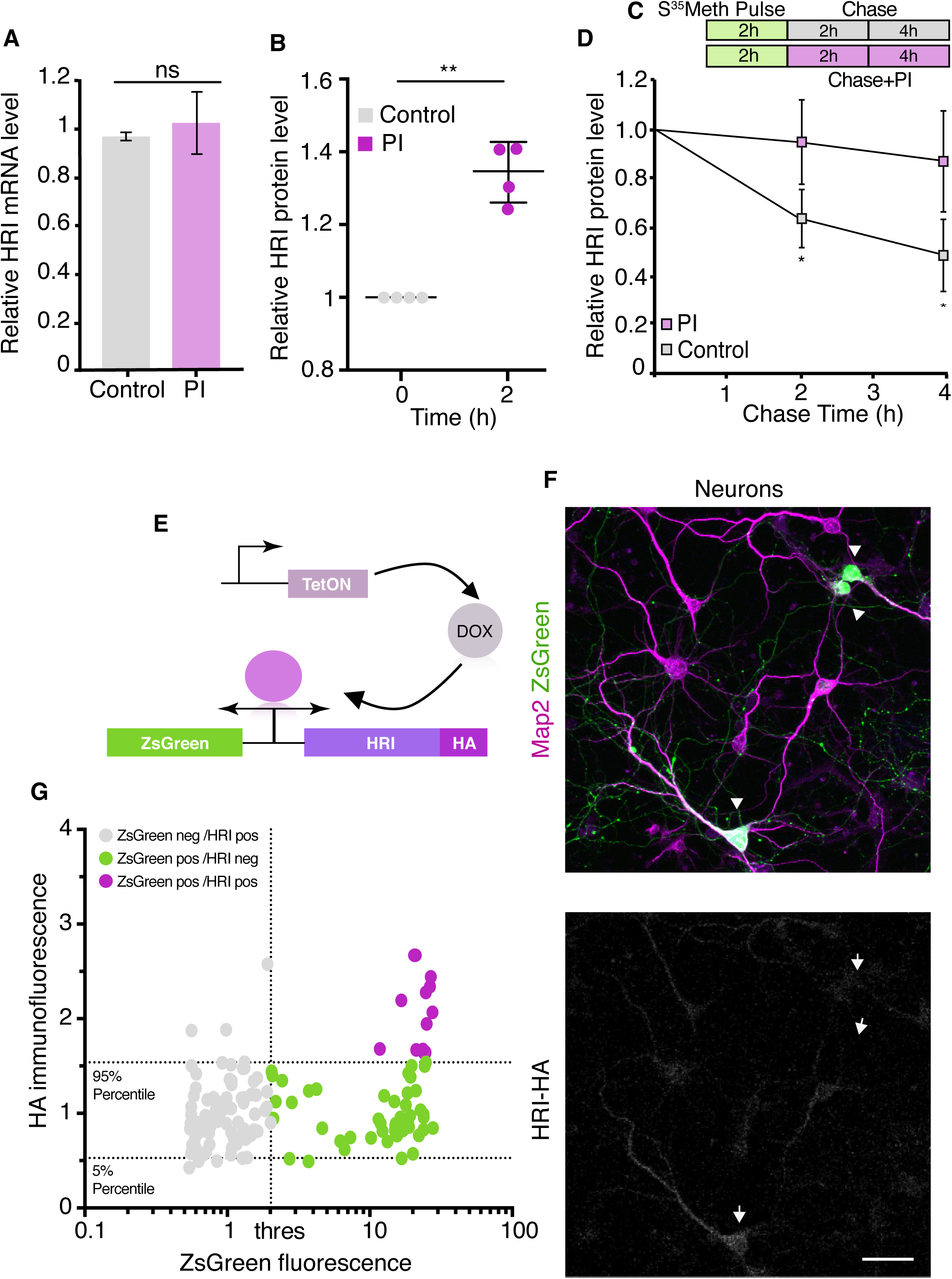
HRI exhibits low expression under basal conditions and increased expression following proteasome inhibition. (A) Analysis for ddPCR experiments showing there is no change in HRI mRNA level following proteasome inhibition, control vs. PI, (unpaired t-test, p >0.05 n = 2, error bars = SD). (B) Quantification of HRI protein levels under control conditions or after PI. After PI (2 hrs) a significant increase in HRI protein was detected in control vs. PI samples, (unpaired t-test p ≤0.01,4 experiments, error bars = SD). (C) Scheme showing the experimental procedure, cultured neurons were labeled with S35-Met for 2h, washed and collected after 2 and 4h +/- PI. (D) Analysis of immunoprecipitated and radiolabeled HRI. Under basal conditions the HRI half-life is ≈4h, its degradation is blocked by PI (unpaired t-test, 2h control vs PI p ≤0.05, 4h control vs PI p ≤0.05, 4 experiments, error bars = SD). (E) Scheme showing the doxycycline-inducible expression of the HA-tagged HRI protein. (F) Representative images of transfected neurons (arrowheads) with the bi-directional reporter resulting in robust expression of ZsGreen (green) but near absent expression of HRI (white) in the same population of cells. Also shown are somata and dendrites (labelled with an anti-MAP2 antibody). Scale bar = 50 μm. (G) Analysis of experiments in F, showing the correlation between ZsGreen fluorescence and HA (HRI) immunolabeling in individual neurons. Dotted horizontal lines indicate the area containing 90% of the HA immunofluorescence values of ZsGreen-negative (non-transfected) neurons (5-95% percentiles), the dotted vertical line indicates the threshold between ZsGreen-negative and ZsGreen positive cells. Grey dots represent ZsGreen-negative cells, magenta dots show neurons positive for ZsGreen but negative for HA, and green dots represent neurons positive for ZsGreen and above the HA 95% percentile of the ZsGreen-negative population (n = 305).

To explore the possibility that there is translational regulation of HRI, we placed both a fluorescent reporter (ZsGreen) and a HA-tagged HRI protein in a single doxycycline (DOX)-inducible bi-directional plasmid (Figure 4E). Transfection of the plasmid into neurons or HeLa cells followed by DOX treatment led to robust expression of ZsGreen in a large population of cells whereas the expression of HRI was absent or barely detectable in the same population of cells (Figures 4F,G and S6). This result was unexpected, given that the use of a bi-directional promoter typically results in roughly equal populations of expressed proteins (Vogl et al., 2018). We next considered the possibility that the mismatch of ZsGreen and HRI-HA expression could be due to the compromised translation of HRI-HA. To address how HRI translation behaves relative to global translation, we used polysome profiling, comparing control and proteasome inhibitor-treated hippocampal cultured neurons. As shown in Figure 5A and consistent with the relative inhibition of translation observed in Figure 1, we observed a leftward shift in the translational profile from polysomes to monosomes following proteasome inhibition. We confirmed this shift using ddPCR to detect beta-actin mRNA levels in each fraction in control and PI conditions (Figure 5B). In contrast, HRI mRNA did not exhibit this leftward shift after proteasome inhibition, but rather exhibited a paradoxical rightward shift to the polysomal fractions (Figure 5C). An Increase in polysome occupancy after proteasome inhibition has also observed for some stress-related transcripts (Thomas and Johannes, 2007) (Figure S7) which exhibit aberrant increases in translation, caused by a variety of manipulations, that are often mediated by uORFs or IRES sequences (Di Prisco et al., 2014; Lang et al., 2002). HRI, however, has a very short 5’UTR and no uORF or IRES is predicted (Di Prisco et al., 2014; Mokrejs et al., 2010; Wu et al., 2009).

**Fig. 5.**
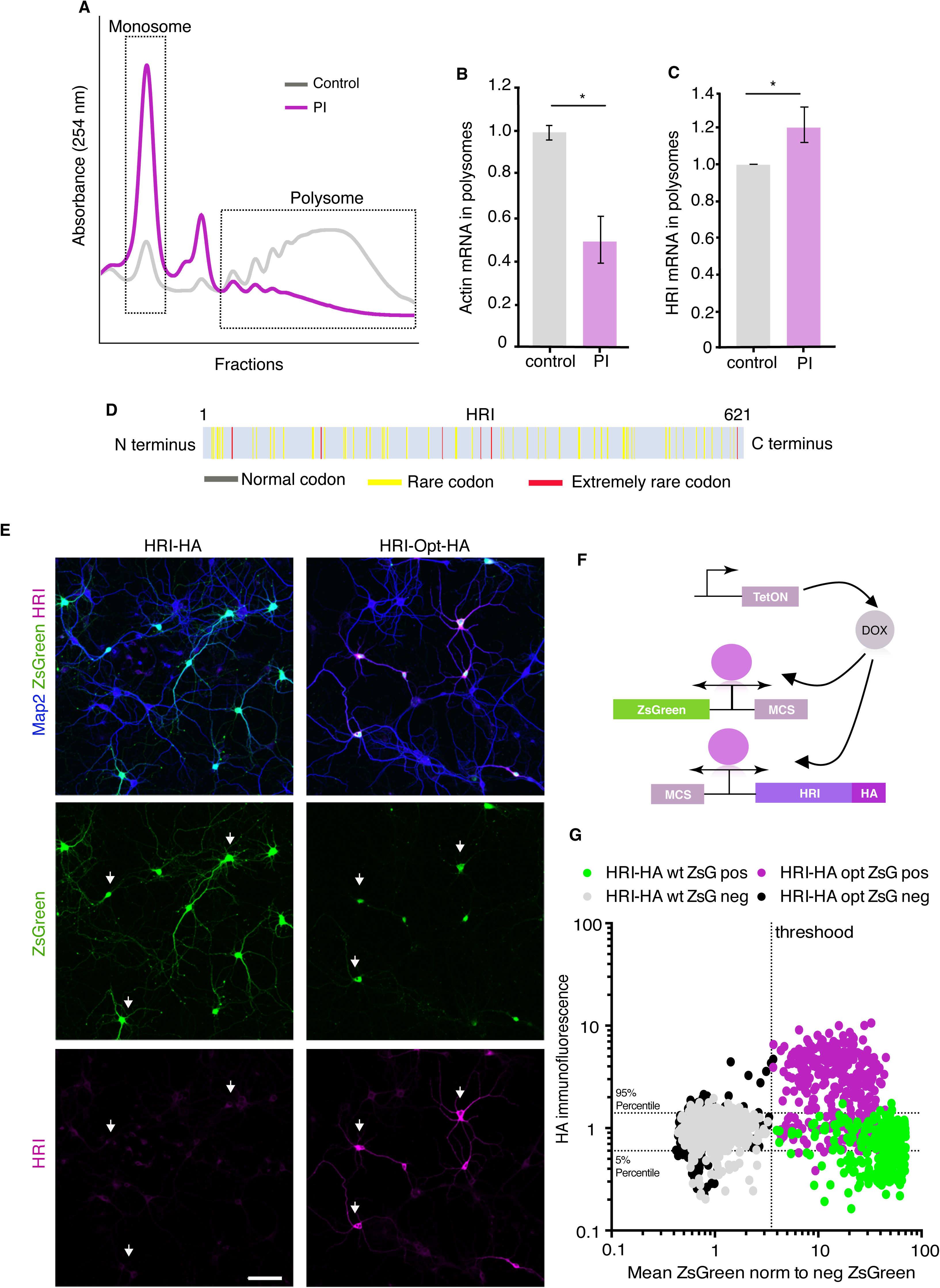
HRI exhibits a codon-dependent paradoxical shift to enhanced translation following proteasome inhibition. (A) Representative polysome profile showing the effect of proteasome inhibition on translation. PI led to a dramatic shift to the monosome fraction, reflecting reduced global translation. (B) Quantification of ddPCR experiments examining the abundance of Actin mRNA in monosomes and polysomes, normalized to control levels. Following PI, actin mRNA exhibited the typical shift to the monosome fraction (unpaired t-test, p ≤0.05, n = 2 experiments (3 technical replicates per experiment). (C) Quantification of ddPCR experiments examining the abundance of HRI mRNA in monosomes and polysomes, normalized to control levels. Unlike the global RNA population, HRI mRNA exhibited a significant shift to the polysome fraction following PI, (unpaired t-test, p ≤0.05, n = 2 experiments 3 technical replicates). Error bars = SD. (D) Scheme of the HRI protein showing an abundance of many rare and extremely rare codons, consistent with HRI’s extremely low level of translation under basal conditions in neurons. HRI uses more rare codons than the 86.78% of the genes expressed in the brain (see Methods). (E) Representative images showing the increased expression of codon-optimized HRI-HA in comparison with HRI-wt (Scale bar = 100μm). (F) Scheme showing the plasmids used for the transfection shown in E, the plasmid shown in the figure 4E was spliced in two parts, one expressing ZsGreen and another one expressing HRI, the plasmids were transfected in 1:5 ratio (ZsGreen:HRI) (G) Analysis of the experiments in E, showing the correlation between ZsGreen fluorescence and HA immunolabeling in individual neurons from dishes co-transfected with ZsGreen and HRI-HA either as wt sequence (HRI-HA) or codon optimized (HRIopt-HA). Dotted horizontal lines indicate the area containing 90% of the HA immunofluorescence values of ZsGreen-negative (non-transfected) neurons of the HRI-HA transfections (5-95% percentiles), the dotted vertical line indicates the threshold between ZsGreen-negative and ZsGreen positive cells. Grey and black dots are represent ZsGreen-negative (non-transfected) neurons from dishes co-transfected with HRI-HA and HRI-opt-HA, respectively. ZsGreen-positive neurons from dishes co-transfected with HRI-HA are shown in green and from dishes co-transfected with HRIopt-HA are represented in magenta. Neurons positive for ZsGreen and HA are located in the upper right quadrant (n=2484 neurons).

An alternative mechanism we considered for the low basal levels of HRI protein is the presence of rare codons in the protein coding sequence, which can result in reduced translational efficiency (Cannarozzi et al., 2010; Chevance et al., 2014; Frumkin et al., 2018; Zhou et al., 2016). We evaluated the rodent HRI sequence for the presence of rare codons and found a number of rare and extremely rare codons, HRI uses more rare codons than 86.78% of the genes expressed in the hippocampus (see Methods), predicting compromised translation (Figure 5D). Indeed, when we expressed an HA-tagged HRI with optimized codon usage (HRI_opt_-HA) we observed that the basal expression of HRI_opt_-HA was higher (Figure 5E,G and S7) and HRI_opt_-HA exhibited a reduced rightward shift to the polysome fraction following treatment with a proteasome inhibitor (Figure S7). Taken together these data indicate that HRI protein expression is paradoxically enhanced under conditions in which global protein synthesis is reduced. Two complementary mechanisms act to enhance HRI levels; first, a stabilization of the protein via reduced degradation by the proteasome and, second, enhanced translation owing to the increased availability of rare codons.

### Stabilized HRI is constitutively active in neurons

In erythrocyte precursors, HRI kinase is activated by multiple mechanisms, including an increase in reactive oxygen species (ROS) or nitric oxide (NO) or reductions in cellular heme (Chen, 2000; Chen and London, 1995; Chen et al., 1991; Igarashi et al., 2004; Uma et al., 2001). Increases in ROS have been reported following proteasome inhibition (Ding et al., 2006). We thus tested whether the HRI activation that follows proteasome inhibition is due to increases in either ROS or NO by quenching ROS or blocking NO synthesis during MG132 treatment. We found that neither ROS scavengers nor NO synthase inhibitors prevented the proteasome inhibitor-induced decreases in protein synthesis (Figure S8). We then examined whether heme metabolism drives the activation of HRI. The major enzymatic path that promotes the breakdown of heme are the heme oxygenases (HO1 and 2). Treatment with two different heme oxygenase inhibitors, however, also failed to affect the protein synthesis reduction elicited by proteasome inhibition (Figure S8). These data suggest that enhanced heme breakdown, *per se*, is not responsible for HRI activation.

In erthryocytes, HRI is activated by reductions in cellular heme levels. We next considered the possibility that basal heme levels might be low in neurons, relative to erythrocytes, which could lead to constitutive kinase activity. We directly measured free heme in neurons and blood and found that while blood had predictably high levels of free heme, heme could not even be detected in cultured hippocampal neurons, and total hippocampal lysates exhibited heme levels orders of magnitude lower than blood (hippocampus; 36±8 fmol/μg, blood; 126±18 pmol/μg). We next tested if the measured amount of heme in blood could indeed inhibit HRI kinase activity in an *in vitro* assay using recombinant HRI and its substrate eIF2a, together with ^32^P-ATP. We found that the addition of hemin (heme complexed with Fe^3+^) led to a significant reduction in eIF2a phosphorylation by HRI (Fig 6A). Given the clear heme-induced inhibition of HRI, we directly tested whether the cytoplasmic regulatory context for HRI is different in neurons by adding recombinant HRI and its substrate eIF2a, together with ^32^P-ATP to cellular lysates prepared from neurons or blood. We evaluated the phosphorylation of eIF2a and found that it was elevated in hippocampus when compared to blood, indicating that HRI is more active under basal conditions in the neuronal cytoplasmic context (Figure 6B). If the elevated HRI protein is constitutively active due to low neuronal heme levels, then the inhibition of protein synthesis induced by proteasome blockade should be amenable to rescue by exogenous heme in neurons. Indeed, we found that addition of hemin to cultured neurons resulted in a rescue of the protein synthesis inhibition caused by proteasome blockade (Figure 6C). Taken together, these data indicate that relatively low cytoplasmic heme levels in neurons favor the constitutive activity of HRI.

**Fig. 6.**
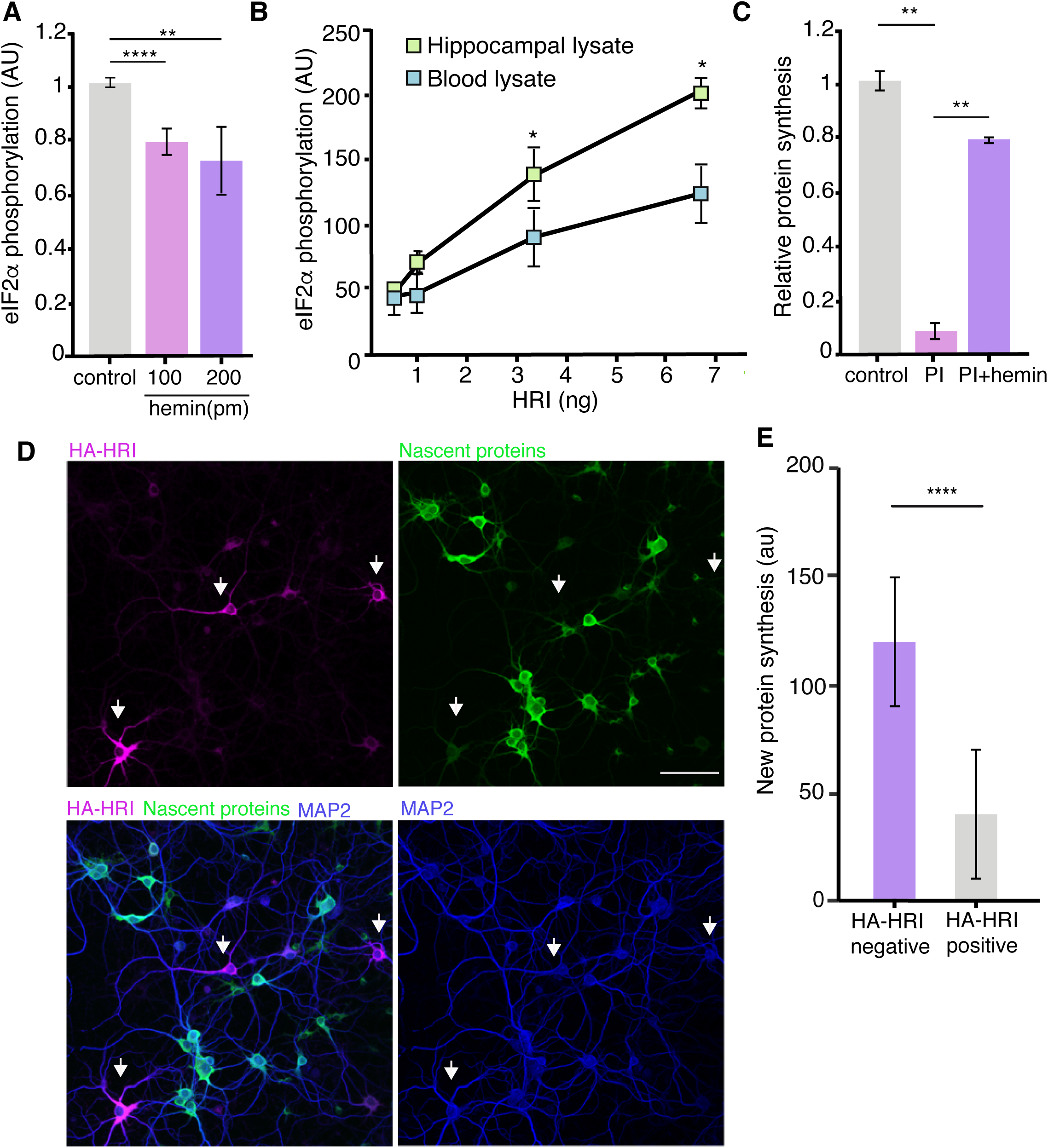
HRI activity is constitutive owing to low heme content of the neurons. (A) In vitro eIF2a phosphorylation assay by HRI; 100 and 200 pm of hemin were added to the reaction mixture, inhibiting the activity of the kinase. (B) In vitro eIF2a phosphorylation assay by HRI; a constant amount of eIF2a was added for each reaction combined with increasing amounts of HRI as indicated (See methods). Increasing amounts of HRI protein lead to significantly higher levels of eIF2a phosphorylation in hippocampal lysates, compared to blood lysates (unpaired t-test 0,05 ng HRI Blood vs hippocampus p ≥0.05, 0,1 ng HRI Blood vs hippocampus p ≥0.05, 0,33 ng HRI Blood vs hippocampus p ≤0.050, 66 ng HRI Blood vs hippocampus p ≤0.05, n=3 experiments). (C) Analysis of experiments testing whether hemin (heme complexed with Fe3+) can mitigate the effects of PI. Hemin significantly reduced the PI-induced protein synthesis inhibition. (PI vs. control, p ≤0.01; PI vs. PI +hemin, Error bars = SD, 3 experiments) (D) Representative images showing that HRI-HA-expressing neurons (arrow) (green; anti-HA labelling) exhibit lower levels of protein synthesis (magenta) when compared to neighboring neurons which do not express HRI-HA. Also shown are somata and dendrites (labelled with an anti-MAP2 antibody) and nuclei (labelled with DAPI). Scale bar = 50 μm. (E) Quantification of somatic protein synthesis levels in neurons expressing or lacking HA-HRI; HA-HRI positive neurons exhibited significantly lower levels of protein synthesis, (p ≤0.0001, HA negative neurons n = 305 and HA positive neurons n=69, unpaired t-test).

The observation that the neuronal cytoplasmic context favors HRI activity suggests that mere expression of HRI could serve as a regulatory switch for protein synthesis. We revisited the HRI over-expression experiments to examine protein synthesis in neurons expressing HRI or not. Brief metabolic labeling of transfected neurons to measure protein synthesis indicated that the small neuronal population that expressed appreciable levels of HRI exhibited significantly reduced protein synthesis, on average, relative to other cells that did not express HRI (Figure 6D, E). This indicates that, in neurons, the cytoplasmic environment is permissive for the constitutive activity of HRI upon expression. In addition, HRI’s potent effect on global translation provides an explanation for why its expression is usually maintained at low levels in neurons. These features of HRI, its basal very low expression, its short half-life and the boost in protein levels upon proteasome inhibition, thus make it both an optimal sensor and effector for proteostasis during situations of compromised proteasome activity.

## Discussion

Here we report that after brief period of proteasomal inhibition (via a pharmacological or genetic manipulation), neuronal protein synthesis is coordinately reduced via the actions of the eIF2a kinase 1, or heme-regulated inhibitor kinase. Taken together, our data indicate the following sequence of events unfolds to restore proteostasis: i) proteasome inhibition (PI) stabilizes the extremely short-lived neuronal HRI protein, ii) endogenous low levels of neuronal heme result in constitutive activation of HRI which increases the phosphorylation eIF2a and reduces translation initiation, iii) the initial decrease in general translation leads to an increase in the availability of rare codons (Saikia et al., 2016), further increasing the translation of HRI mRNA and ensuring ongoing phosphorylation of eIF2a and as a result, reduced global translation.

The above-described mechanism allows for the coordination of protein synthesis and degradation in neurons, so that global protein levels stay balanced. We found that the same feedback mechanism can be detected in isolated neuronal processes, including dendrites and potentially axons. Local synthesis and degradation of proteins occurs in dendrites and axons via localized mRNAs, ribosomes and components of the UPS (e.g. (Bingol and Schuman, 2006; Campbell and Holt, 2001; Hafner et al., 2019; Holt and Schuman, 2013)). It is not known, however, the spatial scale over which this proteostasis operates in axonal or dendritic compartments. In addition, a completely open question in all cells is the how proteostasis is accomplished at the level of individual proteins-of-interest.

Our data indicate that above multiple mechanisms cleverly cooperate to enhance the activity and expression of HRI in response to reduced proteasome activity. These mechanisms make HRI an optimized sensor and effector for sensing and responding to proteasome inhibition. In this regard, it is interesting to note that while the HRI transcript is detected at higher levels than the other 3 eIF2a kinases in hippocampal neurons, the HRI protein is expressed at very low levels under basal conditions, suggesting strong cellular pressure to keep HRI protein expression low. We observed, though, that HRI levels exhibit an increase upon proteasome inhibition, consistent with our observation that HRI can be degraded by the proteasome in *vivo* and *in vitro*. In addition, attempts to over-express HRI via transfection were largely thwarted. We evaluated the transcript for the presence of rare codons which can reduce translational efficiency (e.g. (Frumkin et al., 2018)). Indeed, HRI mRNA possesses many rare and extremely rare codons; expression of a codon-optimized HRI resulted in higher levels of basal translation. In addition, in polysome profiling experiments, we found that HRI mRNA exhibits a paradoxical shift to the polysome fraction following proteasomal inhibition, while the bulk of cellular mRNAs exhibit a shift to the monosome fraction, consistent with their reduced global translation. Owing to the apparent absence of uORFs in the 5’UTR of HRI, we propose that rare codon availability could explain the atypical behavior of HRI mRNA following PI (Saikia et al., 2016), in support of this idea, the codon-optimized HRI failed to exhibit the shift to the polysome fraction upon PI.

In erythrocytes, HRI is activated by reduced heme levels to coordinately reduce the synthesis of α- and β-globin (Han et al., 2001); together α-globin, β-globin, and heme, assembled in a 2:2:4 ratio, make up hemoglobin. Getting the stoichiometry just right is important for red blood cells (RBCs) since an overabundance of any single component is cytotoxic to RBCs and their precursors. We found that heme levels are extremely low or undetectable in neurons, indicating that once expressed at appreciable levels, HRI will be constitutively active. Indeed, we observed significantly elevated HRI activity in a control hippocampal lysate when compared to a control blood lysate. Importantly and definitively, we observed that the PI-induced protein synthesis inhibition was rescued by the addition of hemin.

Taken together, multiple mechanisms act in concert to tightly control the dynamic range of HRI. Under control conditions HRI exhibits barely detectable levels of expression and activity due to its high turn-over and low translation rates. The inhibition of the proteasome is sensed by the stabilization of HRI and the low neuronal heme content means that HRI can respond immediately to the environment created by proteasome blockade. These data further suggest that physiological and pathophysiological conditions that give rise to proteasome dysfunction may favor HRI expression and activity leading to a decrease in protein synthesis.

The role of HRI as a primary mediator of the feedback between the degradation and protein synthesis machinery is surprising given the prominent role the other eIF2a kinases (GCN2, PERK and PKR) are known to play in neuronal translational regulation. Indeed, all four of the eIF2a kinases have been associated with the phosphorylation of eIF2a in response to proteasome inhibition. These studies were performed with cell lines of non-neuronal origin or mouse embryonic fibroblasts (MEF) (Jiang and Wek, 2005; Yerlikaya et al., 2008; Zhang et al., 2010). In addition, the activation of mTOR pathway following UPS inhibition has also been linked to a decrease in protein synthesis (Zhang et al., 2014). In our experiments, HRI is clearly the major kinase responsible for the proteasome inhibition-induced feedback regulation on neuronal protein synthesis; we think it is likely, however, that the other eIF2a kinases also play a role, potentially when HRI is absent or during periods of prolonged proteasome inhibition. Parsing the unique and complementary roles of these kinases will have to take into account the unique translation/degradation rates of each cell type together with the specific expression the eIF2a kinases, protein chaperones and heme content (Uma et al., 1999).

We note that HRI’s role in neuronal function is poorly explored, save for two reports of HRI influencing the translation of synaptic protein (Ramos-Fernandez et al., 2016) and influencing learning (ILL-Raga et al., 2013). In addition, the regulation by heme highlights an important element to consider in the context of neuronal function given emerging data that point to dysregulation of iron homeostasis in disease (Salvador, 2010). Dysfunctional proteasome activity occupies a central position in many neurodegenerative diseases, and increased phosphorylation in eIF2a has been detected in postmortem brains of AD patients (Chang et al., 2002). Prion infection also inhibits proteasomal activity and increases polyubiquitin levels by 2-3 times (similar to the levels observed in our conditions of study- see Fig 1D) (Kang et al., 2004). Strikingly, reperfusion after ischemic stroke leads to a profound proteasome inhibition in brain tissue (Hochrainer et al., 2012). In addition, in hemorrhagic strokes heme is liberated in the brain (Babu et al., 2012; Prabhakaran and Naidech, 2012) and causes neuronal toxicity. The expression of HRI in neurons and its susceptibility to heme regulation should thus be considered in the realms of neurodegeneration and iron dyshomeostasis.

## Materials and Methods

### Primary neuronal cultures

Neuronal cultures were prepared and maintained as described in (Aakalu et al., 2001). Briefly, the cortex from E17 mice or hippocampus from P1 rats were dissected, dissociated with papain (Sigma) and plated on poly-D-lysine coated dishes. Neurons were maintained at 37°C and 5 % CO_2_ in Neurobasal-A plus B27 and GlutaMAX, (Life Technologies). One day after plating neurons were incubated with 3 uM AraC (Sigma) for two days, then the AraC was removed by changing the media. Unless otherwise indicated, neurons were used for experiments at DIV 14-21. For the experiments with KO mice cortical neurons were used, hippocampal neurons where used for the rest of the experiments.

### Transgenic animals

PERK KO (line #009340) and GCN2 KO (line #008240) mice were purchased from the Jackson laboratory. The HRI KO mouse line was kindly provided by Dr. Jane Chen (Han et al., 2001). All lines were maintained as heterozygotes. For neuron cultures, E17 pups from KO/WT x KO/WT breedings were genotyped as indicated by the manufacturer or (Han et al., 2001). Cortices from WT and KO animals were pooled separately and dissociated and plated as described above. Procedures involving animals were performed according to German and Max Planck Society animal care guidelines and approved by and reported to the local governmental supervising authorities (Regierungspräsidium Darmstadt).

### Metabolic labeling, BONCAT and tissue preparation

Neurons were grown and treated with various agents (proteasome inhibitors and other inhibitors) in Neurobasal A (Invitrogen). After treatment, the medium was changed to Neurobasal A lacking Methionine and supplemented with 4 mM AHA +/- treatments for 10 minutes. Neurons were washed 3x with PBS and lysed in PBS with 1% (w/v) Triton X100, 0.4% (w/v) SDS, protease inhibitors w/o EDTA (Calbiochem, 1:750) and benzonase (Sigma, 1:1000), heated at 95°C and centrifuged. BONCAT was performed as described in (Dieterich et al., 2006). In brief, 30ug of total protein was subjected to a click reaction with 300μM Triazol (Sigma, ref. 678937), 50 μM biotin-alkyne tag (Thermo, ref. B10185) and 83 μg/mL CuBr at 4°C o.n. in the dark. Biotinylated proteins were detected by Immunoblot. For puromycylation, puromycin was added to the culture media +/- treatments to a final concentration of 1.5 ng/ul for 10 minutes, then the neurons were washed 2x with PBS and lysed. For phosphorylation studies a phosphatase inhibitor cocktail (Invitrogen) was added to the lysis buffer. Tissue from HRI KO or wild-type mice was homogenized in lysis buffer (1% (w/v) Triton X100, 1% (w/v) SDS, protease inhibitors (1:750) and benzonase (1:500), heated to 95°C and centrifuged. SDS-PAGE was performed with 3ug of lysate and proteins were subsequently blotted onto PVDF membranes.

### Metabolic labeling of dendrites and axons (neurites)

1 million dissociated rat hippocampal neurons were plated into 6-well filter inserts with a 3um pore size (Falcon). The day after plating AraC (3uM; Sigma) was added to the cultures and medium was changed two days later. At DIV8 the proteasome inhibitor MG132 (10uM) was added to the medium. After 2h of treatment the upper part of the filter was carefully scraped with PBS to remove the cell bodies. The dendrites and axons remaining on the other side of the filters were washed 3 times in Neurobasal A-lacking Methionine and incubated for 5min in Neurobasal A-lacking Methionine supplemented with 4mM AHA and MG132 (10uM). Dendrites were collected in lysis buffer and BONCAT was performed as described above.

### Cell viability

Neurons were plated in 96mw plates (20.000 neurons per well) and grown in Neurobasal A without phenol red. The viability was measured with a cell counting kit-8 (Sigma) following the manufacturer instructions. Cell counting solution was added to the media for 1h after 2h of PI incubation (3h of PI incubation in total).

### In situ puromycylation, RNA-FISH and immunocytochemistry

In puromycylation experiments, rat hippocampal or mouse cortical neuron cultures grown on MatTek glass bottom dishes were treated ±proteasome inhibitors and other drugs for the indicated times before puromycin was added at a final concentration of 3.0 ng/µl for 5 minutes then cells were washed 2 times with PBS-MC before fixation and immunocytochemistry. In FISH or immunocytochemistry experiments without puromycylation the incubation medium was removed and cells were immediately fixed for 20 min with PBS containing 4%Sucrose and 4%PFA, pH = 7.4. Panomics RNA-FISH was performed as described previously (Cajigas et al., 2012) using the ViewRNA ISH Cell Assay Kit with 550 dye detection (ThermoFisher). For immunocytochemistry, cells were permeabilized for 15 min with 0.5% Triton in blocking buffer (BB) (PBS with 4% goat serum), blocked in BB for 1h and incubated with primary antibodies in BB for 1 h. After washing, secondary antibodies in BB were applied for 30 min followed, when necessary, by a 3 min incubation with 1 ug/ul DAPI in PBS to stain nuclei. Cells were washed and imaged in PBS and mounted with Aquapolymount (Polysciences) for storage.

### Proteasome peptidase activity

Proteasome activity was measured as described (Martin-Clemente et al., 2004). In brief, after treatment with proteasome inhibitors the neurons were washed 2x with cold PBS and lysed (50mM Hepes pH 7.4, 50mM NaCl, 5mM EDTA, 10uM leupeptin, 1 μg/ml pepstatin and 1 mM PMSF). Lysates were homogenized and centrifuged at 3000g for 10 min at 4°C. Supernatants were immediately used for measurement of proteasome peptidase activity by cleavage of the fluorogenic peptide *N*-Suc-LLVY-MCA (Sigma), enzymatic activity were acquired for 2h. Fluorescence was detected in a micro-plate reader (TECAN) with 400/505 filters. To estimate background signal produced from cleavage by other proteases, proteasome inhibitors were added to lysates from untreated controls.

### Antibodies

The following antibodies were used for immunofluorescence (IF) and/or immunoblotting (IB) at the indicated dilutions: Rabbit anti-peIF2a (for PLA 1:6000 #44728G Invitrogen, and IB, 1:1000, Cell Signaling), mouse anti-eIF2a (IB, 1:1000, Cell Signaling). peIF4B, p4EBP1, peIF4G (IB:1:2500, cell signaling). Most of the commercially available antibodies that should recognize rodent HRI failed to robustly detect the protein in neurons. Nevertheless, the use of HRI knock-out tissue allowed us to identify in neurons, after immunoprecipitation, an HRI-specific band only with one HRI antibody: 07-728, Millipore. Rabbit anti-actin (1:5000, Abcam), rabbit anti-H3 (1:10000, Abcam) rabbit anti-biotin (IB, 1:1000), mouse anti-puromycin (IB, 1:1000, IF 1:3500, Kerafast,), guinea pig anti-MAP2 (IF 1:1000, Synaptic Systems), rabbit anti-HA-tag (IF 1:2000, Rockland) Goat anti-mouse or anti-rabbit IR680 or IR800 (IB, 1:5.000, Licor), goat anti-guinea pig Dylight405 (IF 1:1000, Dianova), goat anti-guinea pig-Alexa488, and goat anti mouse-Alexa546 or -Alexa488, goat anti-rabbit Alexa647 or -Alexa546 (IF all 1:1000, ThermoFisher).

### Immunoprecipitation

The anti-HRI antibody was conjugated with Dynabeads (protein G, Invitrogen) o.n. at 4°C. 30 million mouse cortical neurons (DIV10) were used per IP reaction and lysed in 50mM NaCl, 0.5% NP40 buffer. The lysate was centrifuged at 16,000g for 15 minutes. The supernatant was incubated with the antibody-bound beads o.n. at 4°C. After binding, the beads were washed 4x times for 20min with lysis buffer at 4°C. Bound proteins were eluted from the beads with 0.1M glycine pH = 2.5, the eluates neutralized, subjected to SDS-PAGE and transferred to PVDF membranes.

### DNA Constructs

Rat HRI was cloned into the Tet on-off system from Clontech (Tet-On® 3G Bidirectional Inducible Expression System with ZsGreen1) and an HA tag was added. For some experiments (as indicated in the text) ZsGreen sequences were removed from the plasmid (HRI-HA Tet-On minZsGreen) leaving only HRI and HA inserted (constructs synthesized by GenScript). Codon-optimized versions were made in HRI-HA Tet-On minZsGreen plasmids using GenScript algorithms including optimization of the HA tag. HA-Ubiquitin K48R-G76A (construct synthesized by GenScript).

### Transfections

Neurons were transfected with Magnetofectamine (Life Technologies) using 1.5ug of DNA per 40K neurons for 1h in Neurobasal A (Invitrogen). After transfection, the transfection medium was replaced by conditioned medium. When the plasmid HRI-HA Tet-On minZsGreen was used (see constructs above), it was mixed in a 5:1 proportion with a plasmid containing only ZsGreen under the same bidirectional, inducible promoter to control for transfection efficiency (ten times excess of HRI-HA Tet-On minZsGreen). One week after transfection, Doxycycline (1ug/1ul, Invitrogen) was added to the cultures for 12h before puromycylation. HeLa cells (ATCC) were grown in Dulbecco’s modified eagle’s medium (DMEM, Invitrogen) with 10% fetal calf serum (FCS) at 37°C in an atmosphere with 5% CO_2_ and transfected with lipofectamine (Invitrogen) according to the manufacturer’s instructions. Contamination with mycoplasma was evaluated by PCR (eMyco kit, Intron Biotechnology). After 24h of transfection Doxycycline was added and experiments were performed.

### HRI codon usage

A Codon adaptation index (CAI) (Sharp and Li, 1987) was used to calculate the relative abundance of rare codons in HRI compared with both the transcripts expressed in the hippocampus (12643) and 566 kinases expressed in the hippocampus (Newman et al., 2016). HRI has a lower CAI value than the median for all genes (median for Eif2ak1 = 0.7637 vs. all genes= 0.7995 or kinases =0.8005) in both cases CAI value for Eif2ak1 is in the lowest quartile of the observed CAI distributions. Compared to the distributions Eif2ak1 has the following percentile: 1.) all genes => 13.2238 (Eif2ak1 uses more rare codons than 86.78% of the genes expressed in the hippocampus) 2.) all kinases => 9.5456 (Eif2ak1 uses more rare codons than 90.46% of the kinases expressed in the hippocampus). The mouse reference Transcriptome was downloaded from NCBI RefSeq repository (ftp://ftp.ncbi.nlm.nih.gov/refseq/M_musculus/mRNA_Prot/). Gene expression in mouse acute hippocampal slices in control conditions was determined by RNA sequencing experiment previously conducted in the lab (You et al., 2015). The kinase gene list was generated by a keyword search in gene description. Analysis and plotting scripts can be found in the following repository https://github.molgen.mpg.de/MPIBR-Bioinformatics/CodonUsage.

### ddPCR

RNA was isolated from crude lysates or ribosomal fractions with QIAzol, following the manufacturer’s instructions. 2-4ng of RNA were treated with 4 units of DNAse for 20min at 37°C and inactivated for 15min at 75°C. The RNA was converted to cDNA using SuperScript IV (ThermoFisher) with a mixture of Random primers and dT15V primers. 1ul out of 10ul of the resulting cDNA was used for ddPCR (Bio-rad) with Taqman probes (Integrated DNA Technologies; see Table 1 for the sequences) following the manufacturer’s instructions. The specificity of the primers for the HA-tagged versions of HRI was confirmed by ddPCRs in non-transfected neurons. The mRNA counts of each gene were normalized to the rRNA content of the corresponding fraction (ddPCR for rRNA was performed with cDNA diluted 1:10000). The normalized mRNA content of the fractions 7 to 11 (polysome) was related to the fractions 2-4 (monosome) with the following formula: (mRNA polysome)/(mRNA monosome + mRNA polysome) The disome was excluded from the calculations as its role in mRNA translation is not clear.

### In vitro degradation assay

200 ng of recombinant HRI (SignalChem) or 500ng of recombinant eIF2a (abcam) were incubated with 2ug of purified 20S proteasome (Enzo) in 20mM Hepes pH 7.4/2mM EDTA/1mM EGTA for the indicated times at 37°C. When indicated, the 20S proteasome was incubated with 10uM MG132 for 30min prior to its incubation with the kinase. Protein mixtures were loaded onto SDS-PAGE gels and stained o.n. with SYPRO^®^ Ruby Protein Gel Stain as indicated by the vendor. Gels were imaged using Gel Doc™ XR (Bio-Rad). In the figure the relative degradation of HRI+20S to HRI+20S+PI is shown.

### Pharmacological treatments

Proteasome inhibitors MG132 and Lactacystin were purchased from Sigma, dissolved in DMSO and used from 10 to 25 uM as indicated. Porphyrins were dissolved in DMSO and purchased from Millipore (CoPPIX), Sigma (FePPIX or hemin), or Cayman (SnMPIX and ZnPPIX), 4, 6 or 12 uM of each porphyrin was tested. In the figures always the 4uM dosage is shown. Porphyrins were added at the same time as the proteasome inhibitors. NO inhibitors were added 1h before proteasome inhibitors and kept during the proteasome inhibitor treatment. L-NMMA (NG mono methyl L-Arginine, Cayman) was dissolved in water and used at 2 and 6mM. 7-Nitroindazole (7-Ni, Sigma) was dissolved in DMSO and used at 100 and 200uM. N-Omega-Nitro-L-Arginine (L-NNA, Sigma) was dissolved in water and used at 100 and 200uM. S-methyl-L-thiocitrulline (L-SMTC, Sigma) was dissolved in water and used at 10 and 50uM. ROS inhibitors were added 1h before proteasome inhibitor and kept during the proteasome inhibitor treatment; N-acetyl-cysteine (NAC, Sigma) was dissolved in water and used at 1 and 5mM. Ascorbic Acid (AA, Sigma) was dissolved in water and used at 200uM and 500uM. MitoTempo (Sigma) was dissolved in water and used at 10 and 25uM. Trolox (Sigma) was dissolved in DMSO and used at 500uM and 1mM. The PKR inhibitor (Cayman) was dissolved in DMSO and used at a final concentration of 1uM and was added 1h before proteasome inhibitor treatment. Protein synthesis was evaluated for all drugs alone and no differences were observed when compared to the control treated samples (data not shown; except for MG-132 and lactacystin). All experiments and dosages were tested in at least two biological replicates; the presented quantifications are from the higher dosage tested. Actinomycin-D (Sigma) was dissolved in DMSO and used at 10uM/ul. It was added at the same time as the PI when indicated.

### Heme measurements

Hippocampal neurons were grown in 10cm dishes (5 million cells per well), each plate was lysed in 50μl of the buffer kinase buffer (5mM MOPS, 2.5mM β-glycerophosphate, 1mM EGTA, 0,5mM EDTA). The mouse hippocampus was lysed in 200μl of the above buffer. Mouse blood was collected with 3mM EDTA, the sample was centrifuged 5 minutes at 1000 rpm to collect the erythrocytes, the cells were extensively washed in PBS, and lysed with kinase buffer. Equal amounts of protein (measured by BCA assay-Pierce) were used for free heme measurement following the manufacturer’s instructions (Sigma MAK-036).

### HRI activity in the presence of blood and hippocampal lysates

Blood and hippocampal lysates were obtained as indicated above. To inhibit the endogenous kinases and proteases, the lysates were heated at 60C for 10 minutes, and centrifuged, 25 μg of the lysates were added to the kinase reaction mix (per reaction: 0,5 μg of recombinant eIF2a, 0.0005-0.001-0.0033-0.0066 μg of recombinant HRI, and 60μM ATPγ). The reactions were carried out for 45min at 30°C, stopped with SDS-Page running buffer, run on a gel and transferred to a PVDF membrane that was exposed to a phosphorimager screen.

### Imaging immunocytochemistry and FISH

Images were acquired with a LSM780 confocal microscope (Zeiss) using 20x or 40x objectives (Plan Apochromat 20 x/NA 0.8 M27, Plan Apochromat 40 x/NA 1.4 oil DIC M27). Images were acquired in 8 bit mode as Z-stacks with 1024 x 1024 pixels xy resolution through the entire thickness of the cell with optimal overlap of optical z-sections. The detector gain in the signal channel(s) was set to cover the full dynamic range but to avoid saturated pixels. Imaging conditions were held constant within an experiment.

### Image analysis and representation

To quantify puromycin or HA immunoreactivity or ZsGreen fluorescence, three channel maximum intensity projections of the Z-stacks created in the Zeiss Zen software were opened in ImageJ. For single soma analyses all neuronal cell bodies in the image were manually outlined based on Map2 staining (to distinguish from glial cells) and areas saved as ROIs. Hela cell ROIs were outlined based on combined channels selecting a region around the nuclear stain. The image was split into single channel images in ImageJ and the mean grey value for each ROI was measured in the relevant channels. For visualization, maximum intensity projections of the Z-stacks were adjusted for brightness and contrast in ImageJ with the same settings for samples and controls and over the whole presented image. Examples of single dendrites shown were created with the ImageJ Plugin ‘Straighten’ by tracing a dendrite starting from the soma, resulting in left to right in the image representing proximal to distal.

### Polysome profiling

Cytosolic lysates from primary neuronal cultures were prepared as follows: Cell treated with vehicle or proteasome inhibitors were incubated with 100 μg/ml cycloheximide in the medium for 5 min, washed in cold PBS with 100 μg/ml cycloheximide and lysed in 8% glycerol, 20 mM Tris pH 7.5, 150 mM NaCl, 5 mM MgCl_2_, 100 μg/ml cycloheximide, 1 mM DTT, 1% Triton X-100, 24 U/ml TURBO DNase, 200 U/mL RNasin(R) Plus RNase Inhibitor. Lysates were pipetted up and down until homogenization was clear with a 0.4×20mm needle (HSW FINE-JECT) on ice. Lysates were centrifuged for 10 min at 10,000g at 4°C, and the supernatants were used for ribosome fractionation.

For sucrose gradients, all solutions were prepared in gradient buffer (20 mM Tris pH 7.5, 8% glycerol, 150 mM NaCl, 5 mM MgCl_2_, 100 μg/ml cycloheximide, 1 mM DTT). Gradients were prepared by sequentially adding different sucrose concentrations (in order from first added to last, 8 mL of 55%, 0.5 mL of 50%, 0.5 mL of 40%, 0.5 mL of 30%, 0.5 mL of 20%, 0.5 mL of 10%) into the same Thinwall polypropylene tube (Beckman). After the addition of each sucrose solution, tubes were place at −80°C to freeze the content before the next layer. The gradients were stored at −80°C and left for equilibration at 4°C o.n. Then 0.5 to 1.5 OD (measured with NanoDrop at 260nm) of the lysates were loaded on top of the gradients and spun at 36,000 rpm at 4°C for 2h with a SW41-Ti rotor (Beckman). Fractions from each sample were collected every 9 sec using a density gradient fractionation system (Teledyne Isco, intensity used 1), chased by 60 % sucrose 10 % glycerol in water at 850 µL/min, and continuous monitoring at 254 nm using a UA-6 detector. For quantification of monosome and polysomes peaks, the area under the curve was measured using Image J or the following program: https://software.scic.brain.mpg.de/projects/MPIBR/PolysomeProfiler. The same script was used to visualize the overlay of the polysome profiles of different samples.

### Statistics

Statistical significance of the quantification of IF and WB was tested from single or combined experiments (as stated in the figure legends). If multiple experiments were combined all the data are shown in the same plot. When necessary, data were normalized to one condition. Statistical analysis was performed using GraphPad Prism.

