## Supplementary Figures for "Neuronal Proteostasis is mediated by the switch-like expression of Heme-regulated Kinase 1, acting as both a sensor and effector"

**Supplementary Figure 1**

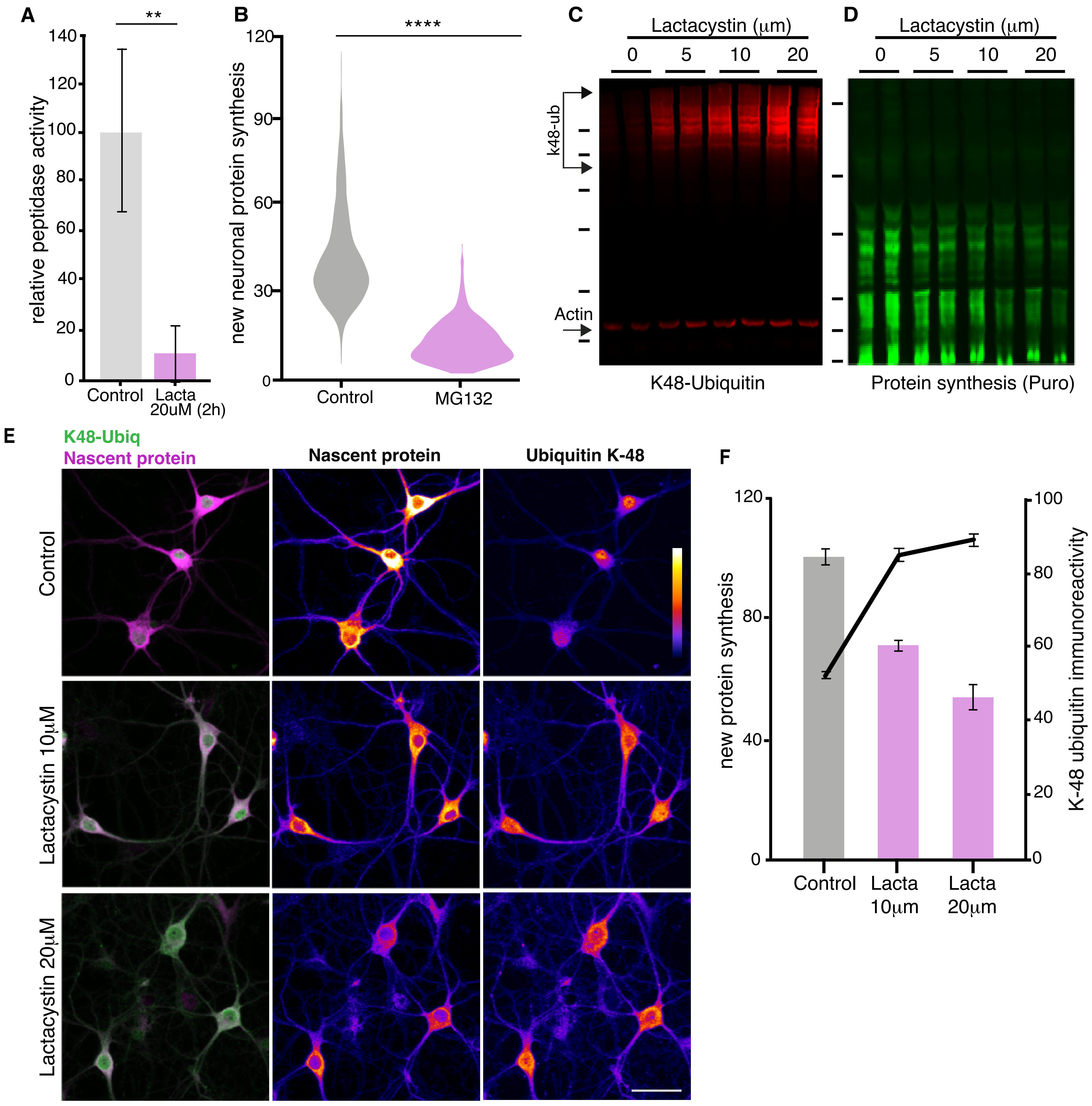


**Fig. S1. Additional experiments examining the effects of proteasomal inhibition on neuronal protein synthesis.** (A) Proteasome peptidase kinetic activity assay showing that a 2 hr Lactacystin treatment inhibits the proteasome by ~ 80% (unpaired t-test, $p \leq 0.001$, control, treated, kinetic slopes are represented, see Methods, 2 experiments). (B) Analysis of experiments shown in Fig 1E. Protein synthesis in hippocampal neurons was reduced by over 30% following proteasome inhibition. ($p < 0.0001$, unpaired t-test, ctrl n= 1768, PI n= 1137 neurons, 5 experiments, 2 dishes per experiment and condition). (C) Representative Western blot of K48 ubiquitin chains. Shown are two biological replicates from a control sample and neurons treated with increasing concentrations of lactacystin (2 hr treatment). Actin is shown as a loading control. Molecular weight markers from top-to-bottom are 250, 150, 100, 75, 50, and 37 kD. (D) Representative Western blot of puromycylated newly synthesized proteins (the same membrane shown in C was incubated with an anti- puromycin antibody). Note that the increasing k48-ubiquitin signal (C) is paralleled by a decrease in puromycin signal. Molecular weight markers from top-to-bottom are 75, 50, 37, 25, 20, 15 kD. (E) Metabolic labeling (using puromycylation, see Methods) of the global nascent protein pool and k48-ubiquitin chains in hippocampal cell bodies and dendrites following proteasome inhibition using increasing concentrations of lactacystin (2 hr treatment). In the first column the merged channels for k48-ubiquitin (green) and puromycin (magenta) are shown, followed by color-lookup images for individual channels. Scale bar = 50 μm. (F) Analysis of experiments shown in E. (unpaired t-test, for protein synthesis; control vs 10μM lactacystin, $p \leq 0.0001$, control vs 20μM lactacystin, $p \leq 0.001$, for k48-ubiquitin; control vs 10μM lactacystin, $p \leq 0.0001$, control vs 20μM lactacystin, $p \leq 0.0001$, control n= 330, lacta 10μM n= 310, lacta 20μM n= 300 neurons, 3 experiments). Error bars = SEM.

**Supplementary Figure 2**

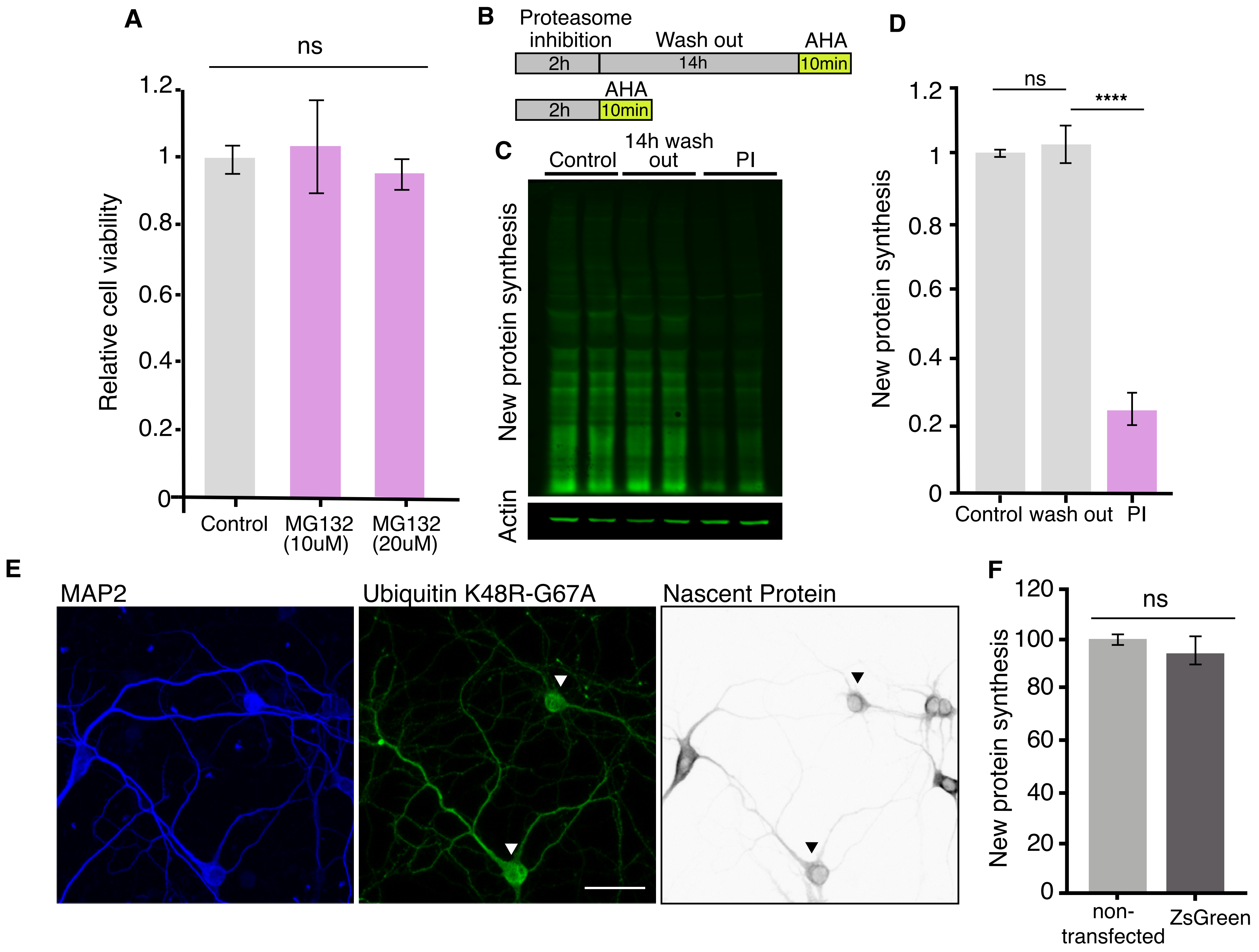


**Fig. S2. Neuronal health assessment after PI, and control experiments for ubiquitin over expression.**
(A) Proteasome inhibition by MG-132 (10 or 20 uM for 2 hr) does not alter cell viability as measured by a cell counting assay (see Methods) (p>0.05, unpaired t-test, control n=20, PI n=20). (B) Scheme indicating experimental protocol: cultured hippocampal neurons were exposed to a proteasome inhibitor (MG132) for 2 hr followed by a 14 hr wash (in the absence of the PI) and then AHA (or puromycin) was added for 10 min to label newly synthesized proteins before or after the washout period (see Methods). (C) Representative puromycylation Western blot showing the recovery of protein synthesis after 2 hr treatment with MG132 and then a 14 hr wash. 2 biological replicates are shown. Actin is shown as a loading control. (D) Analysis of experiment shown in C. Protein synthesis in hippocampal neurons was significantly reduced compared to that observed after the 14 hr wash, which did not differ from control levels of protein synthesis. Control vs. 14 hr wash, not significant, p >0.05; 14 hr wash vs. PI, p ≤0.0001, 4 experiments. All analyses were unpaired t-tests. Error bars = SD. (E) Representative images of transfected neurons overexpressing HA-Ubiq K48R-G76A (arrowheads) (second panel, green), Map2 (blue) is shown in the first panel, nascent protein synthesis (puromycin-labeling) is shown in the third panel (grey), protein synthesis is decreased in the neurons expressing HA-Ubiquitin K48R-G76A compared with non-transfected neurons. Scale bar 50µm. (F) Analysis of the effect that overexpression of a control protein (ZsGreen) has on neuronal protein synthesis. (unpaired t-tests p >0.05, 3 experiments mock n=231, ZsGreen n=316)

**Supplementary Figure 3**

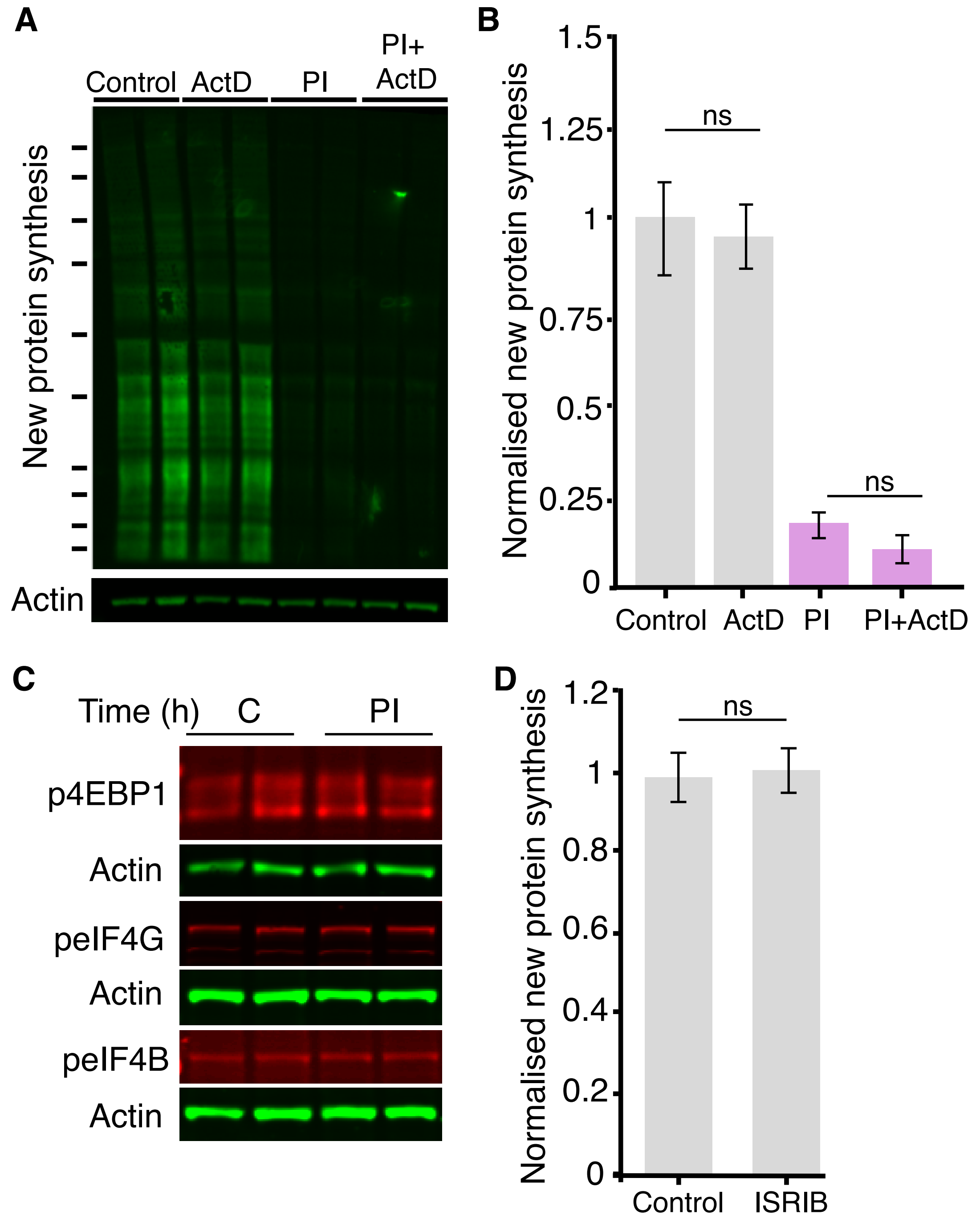


**Fig. S3. Additional data on the eiF2a kinases.** (A) Representative Western blot showing that blocking transcription with actinomycin D (10ug/ul, applied for 2 hr) did not reduce protein synthesis under control conditions or affect the PI-induced decrease in protein synthesis. Actin is shown as a loading control. Two biological replicates per condition are shown. (B) Analysis of experiments shown in A. ActD treatment did not significantly alter protein synthesis under control conditions ($p > 0.05$) nor did it significantly alter the PI-induced decrease in protein synthesis ($p > 0.05$), 3 experiments. (C) Western blot analysis of indicated translation initiation factors and binding proteins showing that proteasome inhibition by MG132 (20µM; 2 hrs) did not alter the phosphorylation levels of 4EBP2, eIF4B or eIF4G. Actin is shown as a loading control. (D) The small molecule inhibitor ISRIB did not significantly alter protein synthesis under basal conditions, control vs. ISRIB, ($p > 0.05$, non-significant, unpaired t-test, 4 experiments).

**Supplementary Figure 4**

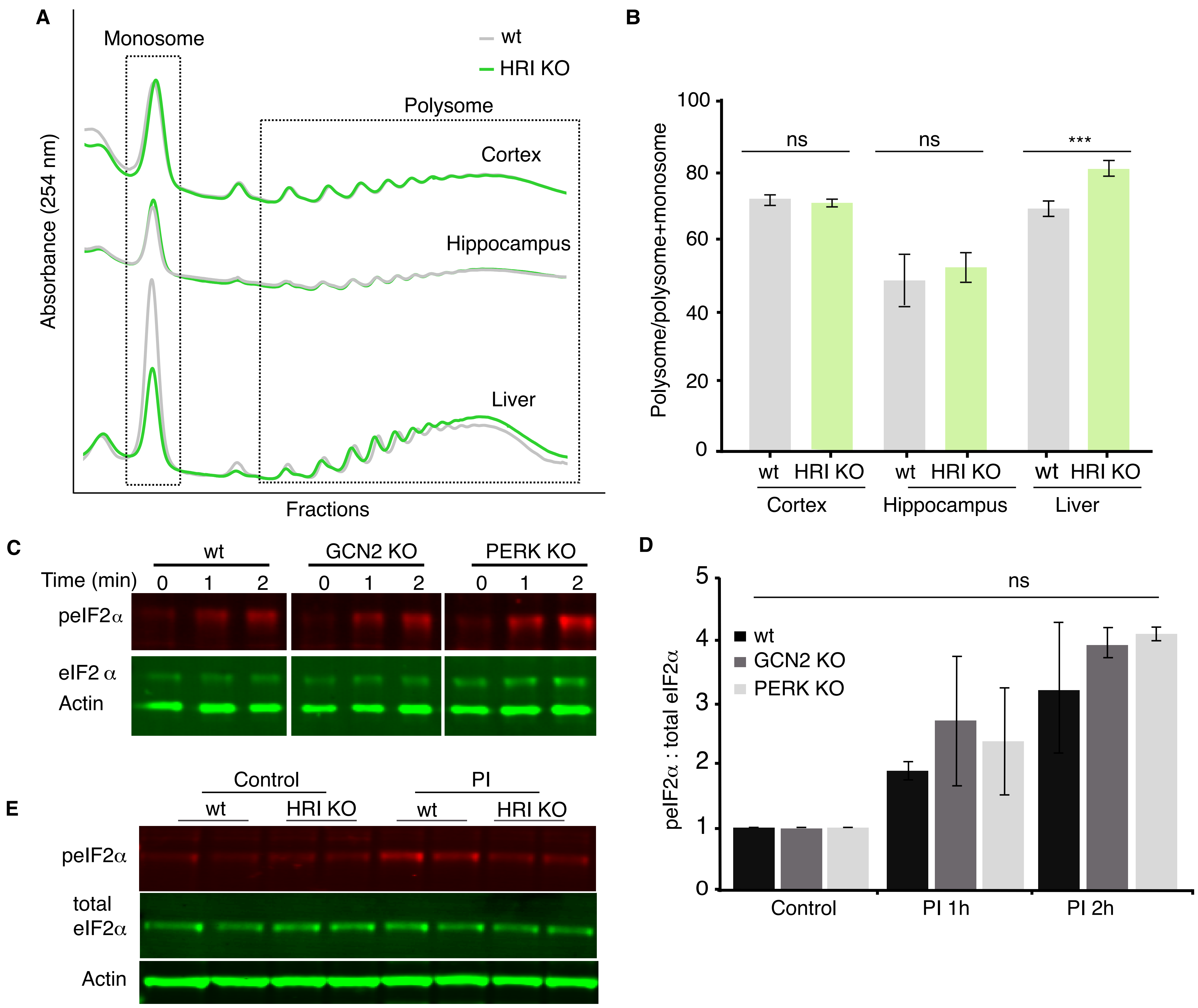


**Fig. S4. Polysome and eiF2a phosphorylation data from eiF2a kinase knock-outs.** (A) Representative polysome profiles from WT or HRI KO cortex, hippocampus or liver. (B) Quantification of the experiment shown in A and additional similar experiments. Shown is the fraction of profiling signal detected in the polysome fraction, expressed as a ration of polysome: polysome + monosome. The absence of HRI (HRI KO) did not change the fraction of signal detected in the polysome in either cortex or hippocampus but did reduce the polysome signal (and enhanced the monosome signal) in liver. WT vs. HRI KO in cortex and hippocampus $p>0.05$, both not significant. WT vs. HRI KO in liver $p \leq 0.001$, n=6 technical replicates and 3 biological replicates for all samples. All analyses used unpaired t-tests. Error bars = SEM. (C) Western blot analysis of eIF2a phosphorylation and total levels following 1 or 2 hrs of proteasome inhibition (MG132, 10 μM) in WT, GCN2 KO or PERK KO cultured neurons. Actin is shown as a loading control. (D) Analysis of experiment in C and additional similar experiments. The PI-dependent increase in peIF2a was not significantly altered in either the GCN KO or the PERK KO. WT vs. GCN2 KO at 1 (n = 2) and 2 hrs (n =2) of PI treatment: $p>0.05$ and $p >0.05$, not significant. WT vs. PERK KO at 1 (n =2) and 2 hrs (n = 2) of PI treatment: $p>0.05$ and $p >0.05$, not significant. All unpaired t-tests, error bars = SD. (E) Representative Western blot showing the levels of peIF2a and total eIF2a following proteasome inhibition in cortical cultures prepared from WT or HRI KO mice.

**Supplementary Figure 5**

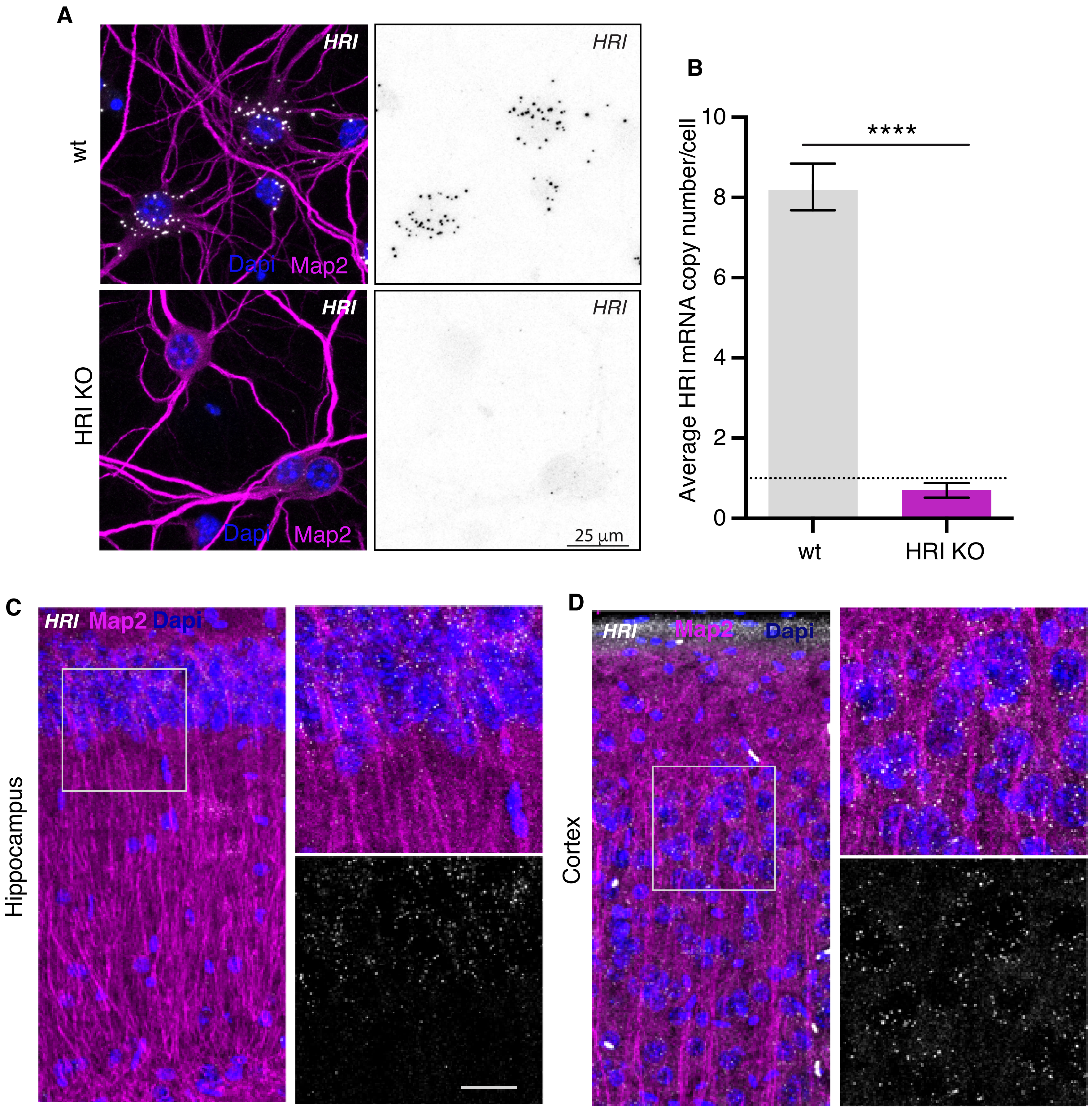


**Fig. S5. HRI mRNA detection by Fluorescence In Situ Hybridization.** (A) Representative fluorescence in situ hybridization images of cultured neurons detecting HRI mRNA in neuronal somata and dendrites of wt and HRI KO mice. HRI mRNA is detected in neurons (white puncta left panel, and black puncta right panel). Scale bar =25 µm. (B) Analysis of the experiment shown in A (wt n=176, KO n=160, 3 experiments). The number of HRI mRNA puncta detected in the HRI KO neurons was significantly reduced relative to control (C) Representative fluorescence in situ hybridization image detecting HRI mRNA (white puncta) in hippocampus and cortex of wt mouse. Scale bar =20 µm. (D) MAP2 (magenta) and Dapi (blue) are shown in all images.

**Supplementary Figure 6**

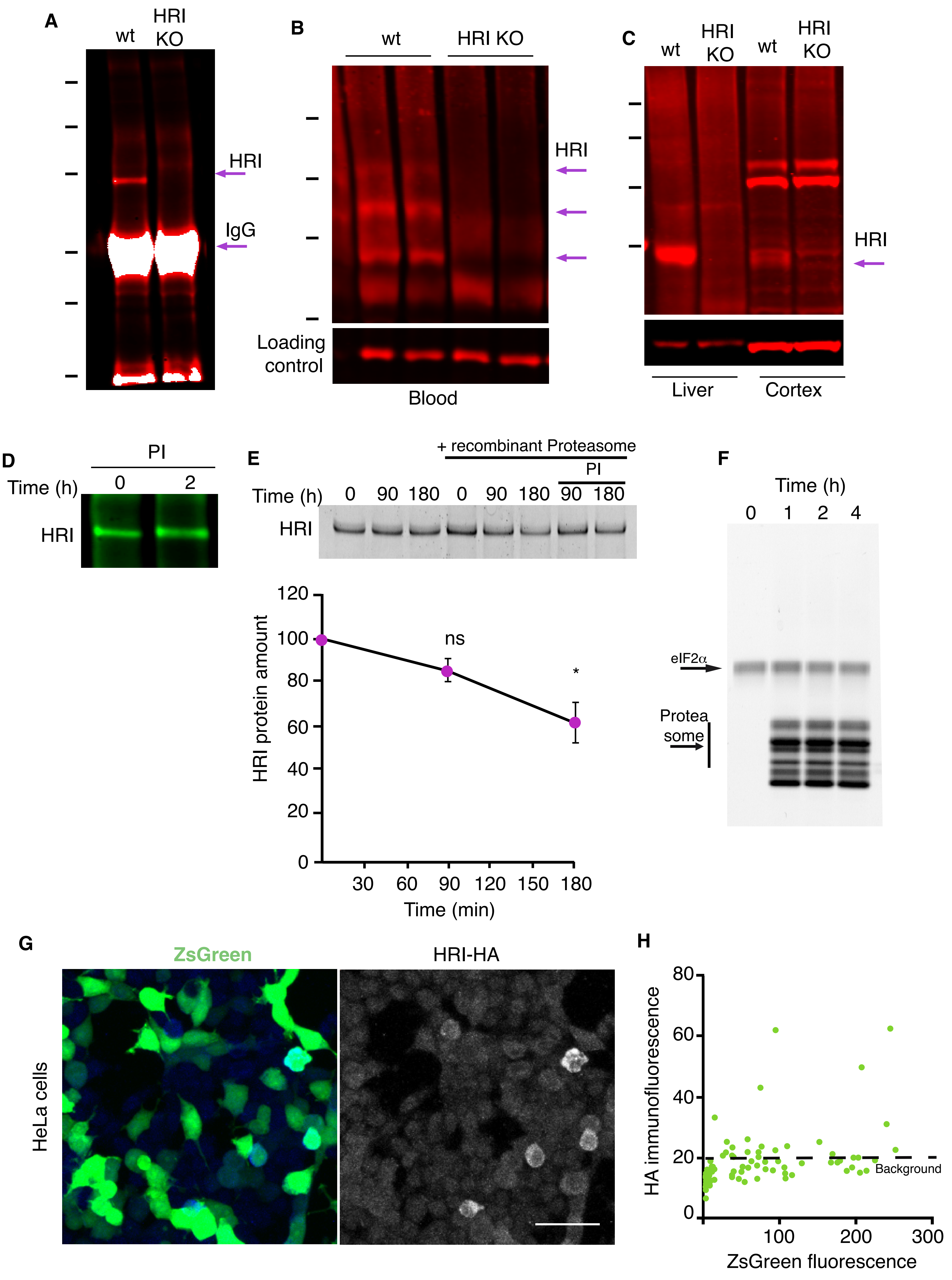


**Fig. S6. Additional data on HRI expression following proteasome inhibition.** (A) Validation of the HRI antibody in cultured neurons showing a distinct band present at 75 kD, (MW of HRI ≈ 75kD) after IP and the absence of the band in cultured neurons prepared from an HRI knock-out mouse. Molecular weight markers from top-to-bottom are 150, 100, 75, 50, 37 and 25 kD. (B) HRI is expressed at significant levels in blood. Arrows point to 3 different apparent HRI variants present in wt and absent in HRI KO. Molecular weight markers from top-to-bottom are 150, 100, 75 and 50 kD. (C) HRI expression in liver and cortex. Molecular weight markers from top-to-bottom are 250, 150, 100, and 75 kD. (D) Western blot demonstrating the modest but significant stimulation of HRI protein expression following a 2hr MG132 (10 µM) treatment. The analysis of this representative experiment and other experiments is shown in Figure 4D. (E) In vitro degradation experiment in which recombinant HRI was added to recombinant 20S proteasome for the indicated times, resulting in the degradation of HRI that was sensitive to proteasome inhibition (PI). Plot shows analysis of the HRI protein degradation by the recombinant 20S proteasome normalized to control reactions without proteasome. Control vs. 90 min, $p > 0.05$, 0 vs 180min $p \leq 0.05$, 2 experiments. Error bars = SD. (F) Control experiment for that shown in (E) showing that eIF2a, as one example, is not a proteasome substrate in this assay. (G) Transfection of the bi-directional reporter in HeLa cells (for scheme see Fig.4E) results in robust expression of ZsGreen but sparse expression of HRI in the same population of cells. Scale bar = 50 µm. (H) Analysis of experiments in G, showing the correlation between ZsGreen fluorescence and HA (HRI) immunolabeling in individual neurons. Dotted horizontal lines indicate the area containing 90% of the HA immunofluorescence values of ZsGreen-negative (non-transfected) cells (5-95% percentiles), the dotted vertical line indicates the threshold between ZsGreen-negative and ZsGreen positive cells. Grey dots represent ZsGreen-negative cells, magenta dots show cells positive for ZsGreen but negative for HA, and green dots represent cells positive for ZsGreen and above the HA 95% percentile of the ZsGreen-negative population Expression of HRI was not positively correlated with ZsGreen expression (3 experiments, n=1105 neurons).

**Supplementary Figure 7**

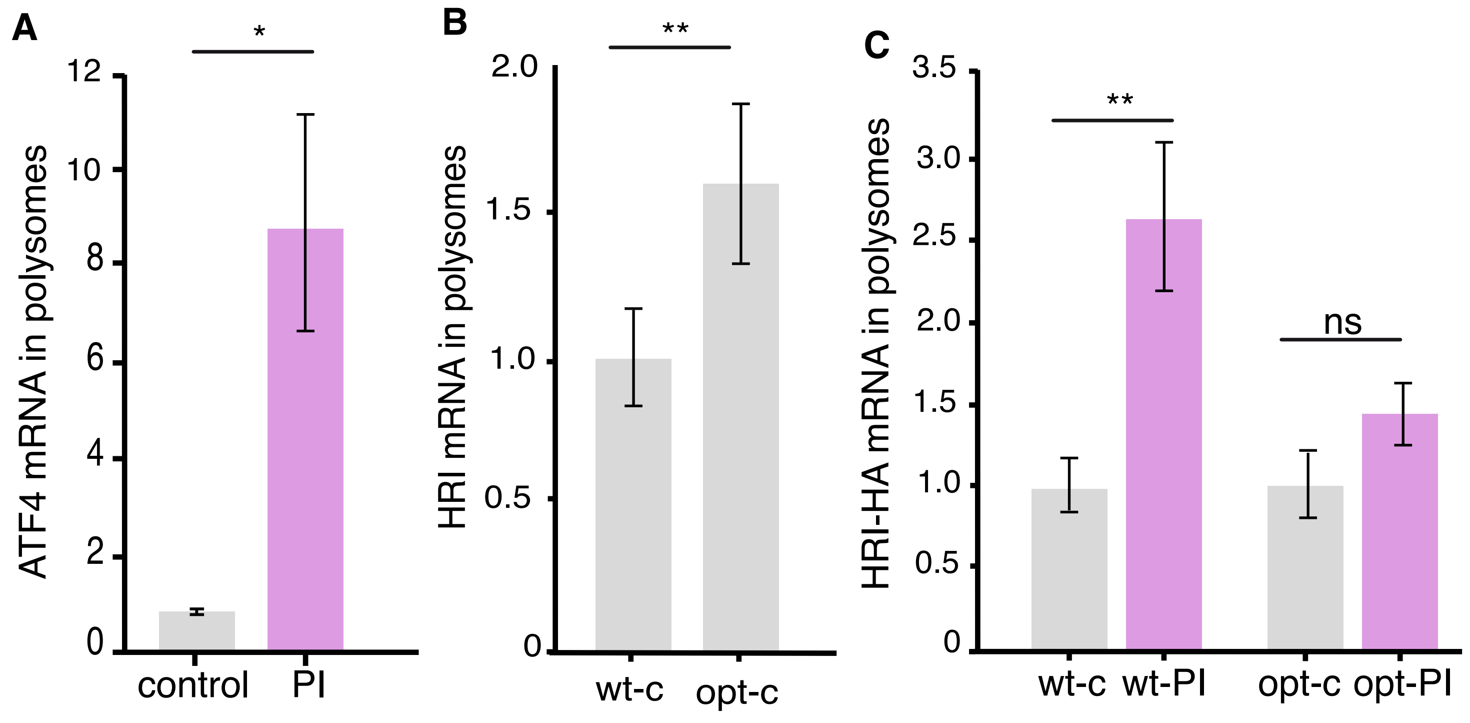


**Fig. S7. Additional data on HRI's codon-dependent paradoxical shift to enhanced translation following proteasome inhibition.**
(A) Control experiments showing the atypical ATF4 mRNA behavior in polysomes after PI (MG132 20 µM, 2 hrs). Following PI, ATF4 mRNA exhibited a noncanonical shift to the polysome fraction, similar to the behavior observed for HRI mRNA (Fig 5C). (Control vs PI $p \leq 0.05$ both unpaired t-test, n= 2 per condition, each replicate has 12 transfected dishes, and 3 technical replicates). Error bars = SD. (B) Analysis of polysome experiment comparing WT HRI (HRIwt) to a codon-optimized HRI (HRIopt). HRIopt exhibited a significantly ($p \leq 0.01$) higher occupancy in polysomes under baseline conditions, indicating enhanced translational efficiency. (n = 2 experiments, each replicate has 12 transfected dishes, and 3 technical replicates). Error bars = SD. (C) Quantification of ddPCR experiments examining the abundance of wild-type HRI (HRI-wt) and a codon-optimized HRI (HRI-opt) mRNA in monosomes and polysomes. Compared to HRI-wt, under control conditions HRI-opt exhibited a higher occupancy in polysomes (B) and upon proteasome inhibition exhibited a blunted shift to the polysome fraction, (unpaired t-test, wt-c vs wt-PI, $p \leq 0.01$, t-test wt-opt vs wt-opt_PI $p>0.05$, n = 2 each replicate has 12 transfected dishes, and 3 technical replicates). Error bars = SD.

**Supplementary Figure 8**

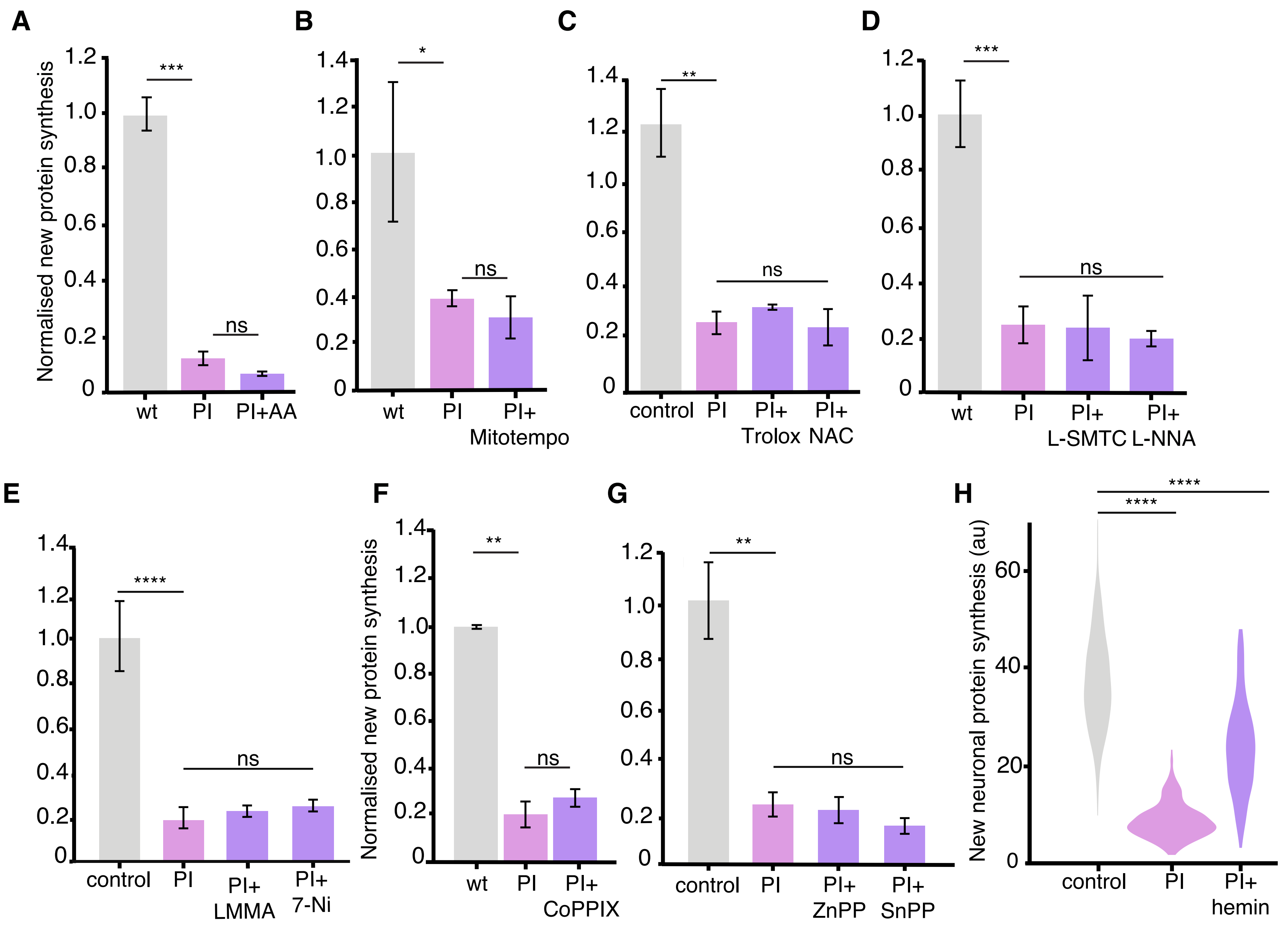


**Fig. S8. Experiments addressing the mechanism of HRI activation.**

Analysis of experiments examining the effect of quenching reactive oxygen species (ROS) (A-C), blocking nitric oxide synthase (D, E) or heme oxygenase (F, G) activity on the PI-induced decrease in protein synthesis (measured using puromycylation- see Methods). None of these treatments resulted a significant alteration of the PI-induced decrease in protein synthesis: in A; PI vs. control, $p \leq 0.001$; PI vs. PI + AA $p > 0.05$; in B PI vs. control, $p \leq 0.05$, PI vs. PI+ Mitotempo $p \geq 0.05$; in C, PI vs. control, $p \leq 0.01$ PI vs. PI+ Trolox and PI+ NAC $p \geq 0.05$; in D PI vs. control, $p \leq 0.001$, PI vs. PI +SMTC and PI +LNNA $p \geq 0.05$; in E PI vs. control, $p \leq 0.0001$ PI vs. PI+ LMMA and PI+ 7-Ni $p \geq 0.05$; in F, PI vs. control, $p \leq 0.01$; PI vs. PI +CoPPIX $p \geq 0.05$; in G PI vs. control, $p \leq 0.01$ PI +ZnPPIX and PI vs. PI+ SnPPIX $p \geq 0.05$ Unpaired t-test for all analyses (n=3 experiments per condition). Error bars = SD. (H) Analysis of experiments examining neuronal protein synthesis in situ using puromycylation showing that hemin significantly rescues the inhibition of protein synthesis in neurons PI (n = 246) vs. control (n = 247), $p \leq 0.0001$; PI vs. PI +hemin (n = 288) $p \leq 0.0001$, Unpaired t-test for all analyses (n=2 experiments, 2 dishes were imaged per experiment) per condition.
